# Uncovering hidden members and functions of the soil microbiome using *de novo* metaproteomics

**DOI:** 10.1101/428334

**Authors:** Joon-Yong Lee, Hugh D. Mitchell, Meagan C. Burnet, Ruonan Wu, Sarah C. Jenson, Eric D. Merkley, Ernesto S. Nakayasu, Carrie D. Nicora, Janet K. Jansson, Kristin E. Burnum-Johnson, Samuel H. Payne

## Abstract

Metaproteomics has been increasingly utilized for high-throughput molecular characterization in complex environments and has been demonstrated to provide insights into microbial composition and functional roles in soil systems. Despite its potential for the study of microbiomes, significant challenges remain in data analysis, including the creation of a sample-specific protein sequence database as the taxonomic composition of soil is often unknown. Almost all metaproteome analysis tools require this database and their accuracy and sensitivity suffer when the database is incomplete or contains extraneous sequences from organisms which are not present. Here, we leverage a *de novo* peptide sequencing approach to identify sample composition directly from metaproteomic data. First, we created a deep learning model, Kaiko, to predict the peptide sequences from mass spectrometry data, and trained it on 5 million peptide-spectrum matches from 55 phylogenetically diverse bacteria. After training, Kaiko successfully identified unsequenced soil isolates directly from proteomics data. Finally, we created a pipeline for metaproteome database generation using Kaiko. We tested the pipeline on native soils collected in Kansas, showing that the *de novo* sequencing model can be employed to construct the sample-specific protein database instead of relying on (un)matched metagenomes. Our pipeline identified all highly abundant taxa from 16S ribosomal RNA sequencing of the soil samples and also uncovered several additional species which were strongly represented only in proteomic data. Our pipeline offers an alternative and complementary method for metaproteomic data analysis by creating a protein database directly from proteomic data, thus removing the need for metagenomic sequencing.

**Significance Statement:** Proteomic characterization of environmental samples, or metaproteomics, reveals microbial activity critical to our understanding of climate, nutrient cycling and human health. Metaproteomic samples originate from diverse environs, such as soil and oceans. One option for data analysis is a *de novo* interpretation of the mass spectra. Unfortunately, the current generation of *de novo* algorithms were primarily trained on data originating from human proteins. Therefore, these algorithms struggle with data from environmental samples, limiting our ability to analyze metaproteomics data. To address this challenge, we trained a new algorithm with data from dozens of diverse environmental bacteria and achieved significant improvements in accuracy across a broad range of organisms. This generality opens proteomics to the world of natural isolates and microbiomes.

## Introduction

The soil microbiome is responsible for carrying out many functions that are important on a global scale, including cycling of carbon and other nutrients and support of plant growth. Over the last few decades high throughput sequencing technologies have made great strides in revealing the soil microbial community composition in a variety of soil habitats and how those communities are impacted by environmental change. Amplicon sequencing has revealed that soil and sediment microorganisms have a very high diversity; much more so than other ecosystems^1^. In addition, metagenome sequencing has proven to be an extremely useful tool for not only determining the composition of soil microbiomes, but also their putative functions. However, not all genes detected in a metagenome survey are actively expressed and significant challenges remain in understanding the biological functions that are carried out by active members of the soil microbiome. Other meta-omics technologies, such as metatranscriptomics and metaproteomics, have helped to close this current knowledge gap. Metatranscriptomics provides information on community transcription and is often used as a proxy for assigning metabolically active members of a soil microbiome. However, metatranscriptomics can only provide a snapshot of gene expression at the moment of sampling. A significant amount of post-transcriptional regulation affects protein abundance and activity^2^. Therefore, metaproteomics provides an essential layer of information about microbiome activity by revealing which proteins are actually produced and have passed transcriptional and translational regulation points.

Despite the promise of metaproteomics for elucidating functions of elusive soil microorganisms, significant challenges remain. An important assumption in most mass spectrometry proteomics identification algorithms is that the set of potential proteins is known, and thus a database of these protein sequences is a typical requirement^3,4^. In environmental samples, however, obtaining an accurate catalog of organisms and their proteins is a challenge, as it is not possible to know the organisms present in the sample beforehand. Amplicon and metagenome sequencing of a matched sample is often used to identify community membership; however, many species might not be observable by sequencing^5–10^.

Here, we present a new method to generate a protein database directly from metaproteomic data as an alternative and orthogonal method of identifying soil microbe composition. The method starts with analyzing mass spectra *de novo* (without a database)^11^, identifying species from the observed peptides and then gathering full proteomic databases for these species. As currently available software tools for *de novo* identification were not sufficiently accurate for environmental samples, we first trained a new deep learning model on spectra from 55 bacteria in nine phyla. After confirming that the new model could successfully identify natural soil isolates, we applied the model to metaproteomics samples. Using a metaproteomics dataset from Kansas soil, our pipeline identified all abundant taxa identified in traditional 16S data as well as identifying new abundant organisms in the soil. Using the identified organisms, we re-analyzed the metaproteomics data and identified differential metabolic functions between species in the microbiome.

## Results

### A new model for *de novo* MS/MS identification

Using a large and environmentally diverse set of mass spectrometry proteomics data, we sought to improve on peptide/spectrum identification where no protein sequence database is available. We adapted a deep neural network structure^12^ and trained a new model called Kaiko, after the Japanese deep ocean submersible used to explore the Marianas Trench. For training and validation, we used 4,604,540 spectra and 927,316 peptides from 51 distinct bacteria (Fig. 1A, Supplementary Table 1). Deep neural networks, like Kaiko, require very large training datasets for parameter optimization. For our neural network architecture, training events with less than 3 million spectra resulted in severely overfit models (Supplementary Figs. 1 and 2). After training and optimization, we evaluated the accuracy of Kaiko against spectra in the test dataset consisting of spectra from four additional organisms not used in model training (511,765 spectra and 90,048 peptides). Kaiko achieved an average accuracy of 33% over all testing files and organisms, a significant improvement over other *de novo* algorithms (Fig. 1B). When considering the top five spectrum annotations, average accuracy exceeded 41%.

**Figure 1.**
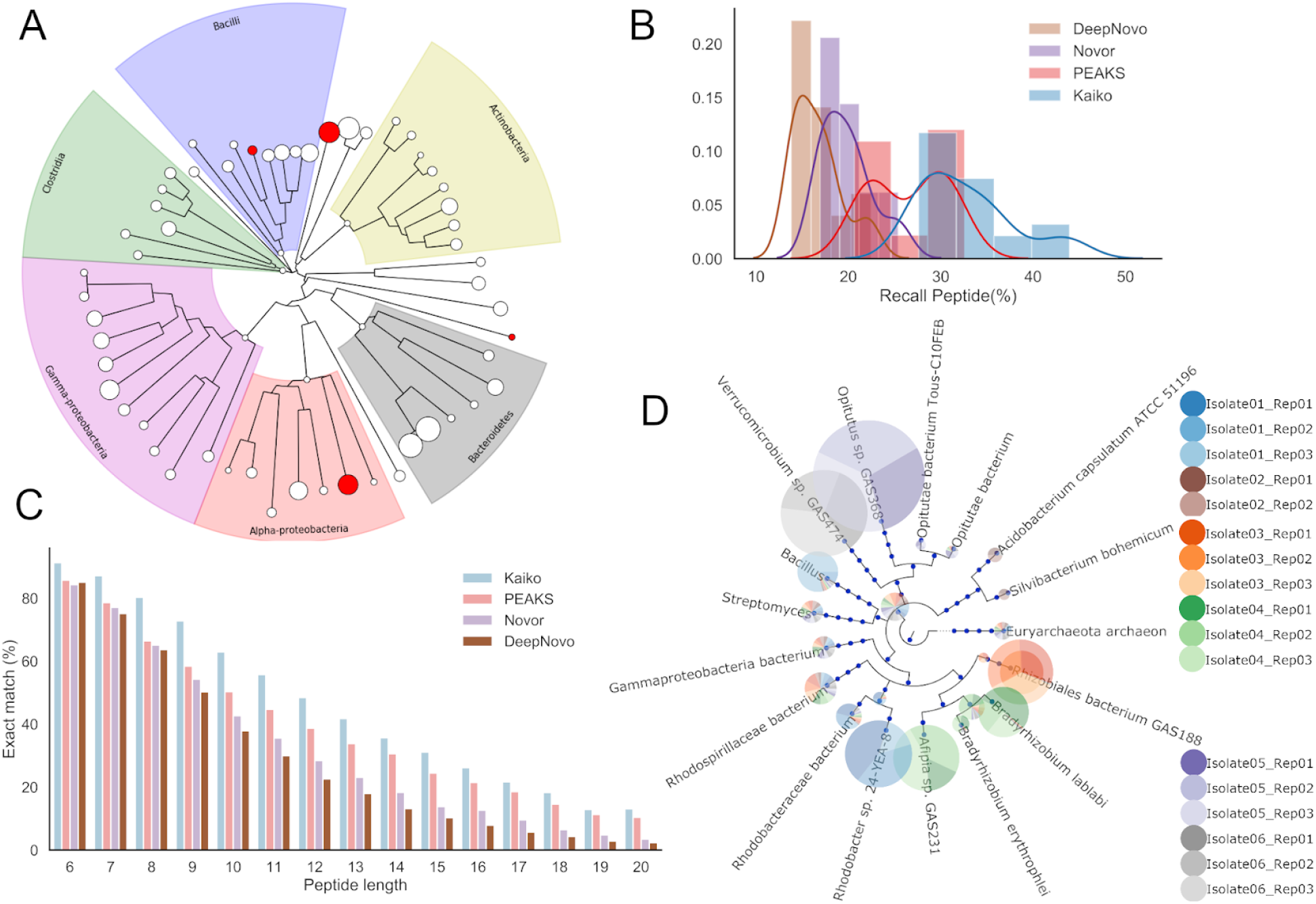
Training, validation and testing of a new *de novo* peptide identification algorithm. (A) Bacteria represented in training and testing data and shown in a phylogenetic tree built from the multiple sequence alignment of rplB is shown for all organisms in the training (white nodes) and testing datasets (red nodes). The size of the node is scaled to represent the number of spectra used. (B) Accuracy of spectrum annotation for four de novo spectrum annotation tools. (C) For each peptide sequence length, the accuracy of spectrum annotation is shown for each of the four algorithms. (D) For each of the six natural isolates, replicate proteomics data was annotated with Kaiko and identified peptides are visualized on a phylogenetic tree. The size of the pie wedge is scaled to represent the number of spectra matching that taxon. For each sample, the top 5 taxa according to the number of peptide hits was included in the visualization.

We next looked at model performance as a function of peptide length (Fig. 1C). Most algorithms performed well with short peptides, length < 8. Unfortunately, these peptides are infrequent in bottom-up proteomics data samples (Supplementary Fig 3). Kaiko exhibited significantly improved accuracy at all lengths, but especially for the most common peptide lengths (10-15 residues), where it achieved an accuracy of ~30-60%. We note that Kaiko had high accuracy at very long peptide lengths of 15 and above. Although these peptides are extremely difficult to annotate *de novo*, they are valuable for predicting phylogeny as the long sequences are more likely to be uniquely mapped to a small taxonomy range.

### Identification of soil isolates via proteomics

Proteomics analysis of natural bacterial isolates from soil often requires *de novo* spectrum annotation. To demonstrate the ability of our deep learning-based algorithm to annotate spectra from an unknown organism and also to accurately identify the unknown organism, we obtained bottom-up proteomics data from six microbes isolated from soil and attempted to identify the sample. For each sample, we annotated the proteomics data with Kaiko and used DIAMOND^13^ to identify the closest sequences in the UniProt database^14^ (see Methods). We then plotted the organisms which had the most matching spectra and inferred the organism for the sample.

For four samples, a matched proteome database became public during our investigation; however, this was still blinded from our analysis. In each of these cases, we identified the exact species as the source of the sample (Fig. 1D). This included two Verrucomicrobia for which Kaiko’s training data had nothing in the same phylum: *Opitutus* sp. GAS368 and *Verrucomicrobium* sp. GAS474. The other two isolates with a matched genome were from the order Rhizobiales: *Afipia* sp. GAS231 and *Rhizobiales* bacterium GAS188. The *Afipia* sample also contained spectra which mapped to neighboring *Bradyrhizobium* species, which could be from shared gene content, contamination or previously unidentified co-culturing.

For two samples, there were no matched proteomes in UniProt and we attempted to derive the true sample identity by 16S sequencing. Isolate 02 cannot be definitively assigned to a genus within NCBI’s taxonomy based on 16S sequencing, but is close to multiple genera within the family Acidobacteriaceae. Using Kaiko’s peptide annotations, we identified two potential candidates for the sample: *Acidobacterium capsulatum* and *Silvibacterium bohemicum* (both Acidobacteriaceae). However, both species had significantly fewer peptide hits matching their proteome and therefore, were weaker matches than expected. This weak alignment to a single organism and splitting between organisms within the same family is consistent with the isolate’s ambiguous taxonomic assignment. The final sample, Isolate 01, was suggested to be a *Gemmobacter* species by 16S sequencing. Peptide hits from Kaiko identified this sample as *Rhodobacter* sp. 24-YEA-8, which is within the same family as *Gemmobacter* (Rhodobacteraceae). With the difficulties surrounding bacterial taxonomic classification and the uncertainty of species designation, this is still a close match.

### Building a protein database without metagenomics

In metaproteomics data analysis, constructing a protein sequence database is a critical component for protein identification^15,16^, as identification sensitivity suffers as database size increases^17^. It is therefore essential to identify organisms present in a sample with taxonomic precision, so that databases include as few species as possible. We present a new solution that derives the organisms present in a sample directly from Kaiko’s analysis of metaproteomics data, thus enabling metagenome-free peptide identification (Fig. 2).

**Figure 2.**
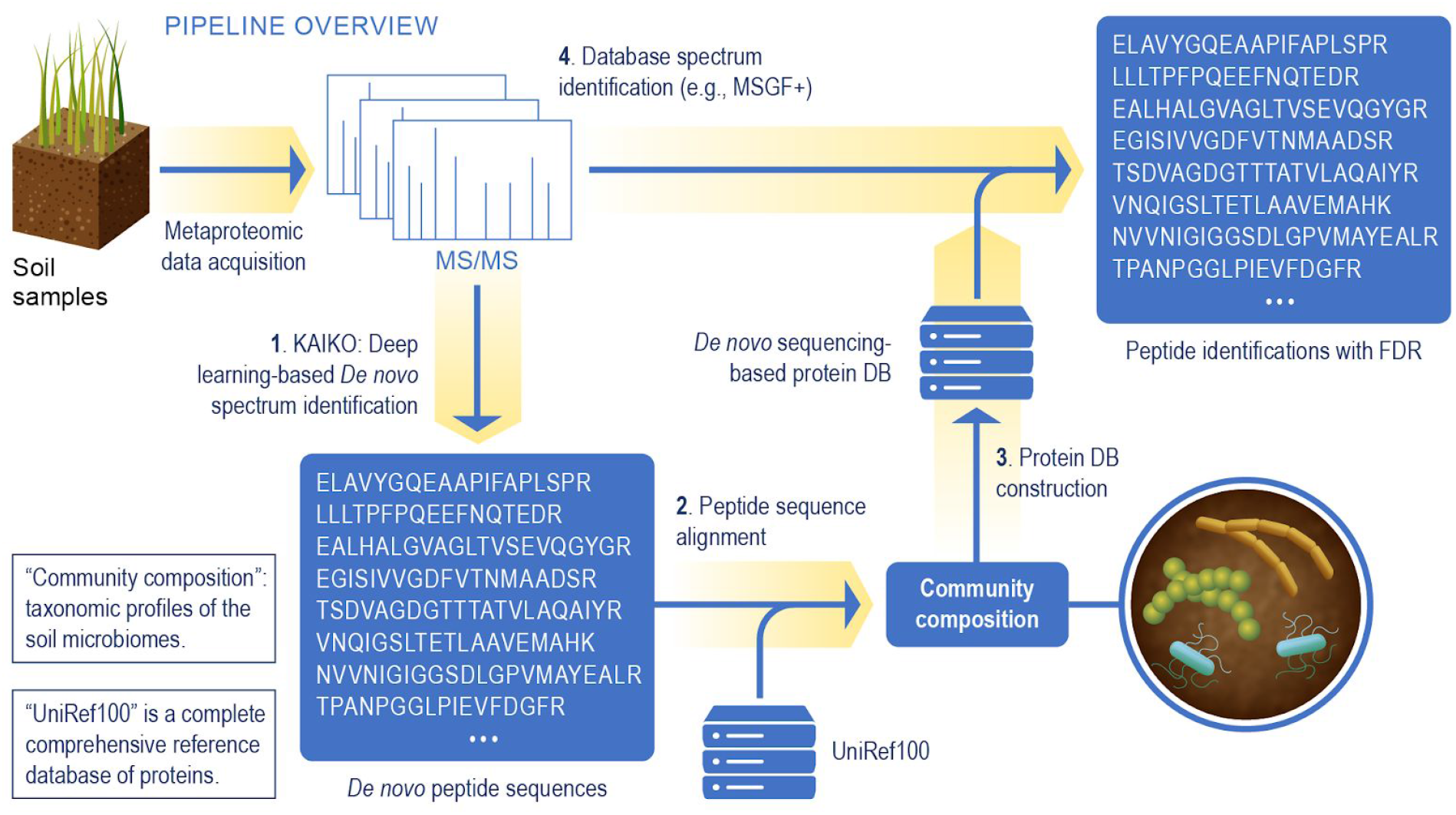
Overview of the metaproteomics data analysis leveraging *de novo* spectrum identification based on the Kaiko model. Peptides are identified using Kaiko, and used to infer community composition (steps 1-3). In step 4, the spectra are re-analyzed using a database search algorithm, e.g. MSGF+, and the protein sequence database created in step 3. This yields a final list of peptide identifications which can be used for functional analysis.

To demonstrate this *de novo*-based metaproteomics pipeline, we analyzed metaproteomic data acquired from pooled samples of native soils collected in three sites located in Kansas^18,19^. To identify species, the Kaiko model and DIAMOND were employed to determine the most dominant organisms, and whole proteomes were retrieved from UniProt (See Methods). 6,410 unique taxa IDs were identified in total and 224 taxa had more than 5 matched peptides. These taxa included well-known bacterial phylotypes consistently detected as a core component of soil ecosystem such as Proteobacteria, Actinobacteria, Acidobacteria, Planctomycetes, Chloroflexi, Verrucomicrobia, Bacteroidetes, Gemmatimonadetes, Firmicutes and Armatimonadetes^20,21^. In addition, our pipeline revealed globally abundant fungal classes such as Agaricomycetes, Sordariomycetes, Eurotiomycetes, Leotiomycetes and Mortierellomycetes^21,22^.

To evaluate the taxa annotation from the Kaiko model, we also identified taxa using 16S rRNA data from the same samples (See Methods). 243 unique taxa IDs were determined for 3,693 OTUs. All of the highly abundant phyla detected by 16S were also detected by Kaiko (Table 1). Several phyla uniquely found by Kaiko are known to be present in environmental soils^23–28^. For example, Candidatus Rokubacteria is distributed globally in diverse terrestrial ecosystems, including soils and the rhizosphere^23^ and Candidatus Tectomicrobia has also been detected in soils^24^.

**Table 1.** Relative abundance of the top 20 bacterial phyla detected from 16S and Kaiko. A dash in the table represents the corresponding phylum was ‘not detected’. The asterisk(*) Indicates that some taxa in the corresponding phyla are used to construct the protein DB.

| Phylum | Read counts<br>By 16S | Peptide<br>counts By<br>Kaiko | Relative read counts<br>% total reads at the<br>phylum level | Relative Peptide counts<br>% By Kaiko at the<br>phylum level |
| --- | --- | --- | --- | --- |
| Proteobacteria* | 40778 | 4903 | 34.6 | 38.1 |
| Actinobacteria* | 16501 | 3949 | 14.0 | 30.7 |
| Acidobacteria* | 18562 | 1010 | 15.7 | 7.8 |
| Firmicutes* | 6761 | 634 | 5.7 | 4.9 |
| Chloroflexi* | 767 | 479 | 0.7 | 3.7 |
| Bacteroidetes* | 9712 | 467 | 8.2 | 3.6 |
| Planctomycetes* | 11427 | 321 | 9.7 | 2.5 |
| **Candidatus Rokubacteria\*** | - | 266 | - | 2.1 |
| Verrucomicrobia* | 11841 | 237 | 10.0 | 1.8 |
| Cyanobacteria | 489 | 162 | 0.4 | 1.3 |
| Gemmatimonadetes* | 869 | 61 | 0.7 | 0.5 |
| Nitrospirae* | 18 | 44 | - | 0.3 |
| **Candidatus Tectomicrobia\*** | - | 43 | - | 0.3 |
| **Deinococcus-Thermus** | - | 32 | - | 0.2 |
| **Spirochaetes** | - | 32 | - | 0.2 |
| **Elusimicrobia** | - | 15 | - | 0.1 |
| Tenericutes | 99 | 15 | 0.1 | 0.1 |
| Armatimonadetes | 75 | 13 | 0.1 | 0.1 |
| Ignavibacteriae | 16 | 6 | 0.01 | 0.05 |
| Chlamydiae | 2 | 4 | 0.00 | 0.03 |

To construct the protein database from the identified organisms, we selected the 100 most abundant bacterial taxa, resulting in a protein database containing 17,448,135 protein clusters (UniRef sequences) from 12 bacterial phyla. We note that the 100 taxa identified by proteome data consist of 91 species, 1 genus, 7 strains, and the remaining 1 had no phylogenetic rank. Unfortunately, the 16S taxa annotations were often resolved only to a phylum or class level; relatively few taxa from 16S data were able to be narrowly identified at the level of genus or species. Creating a protein sequence database for all species within a broad taxonomic category, such as phylum, would dramatically increase the size of the protein sequence database and reduce the sensitivity of the proteomics data analysis.

### Soil metaproteomic data analysis

Using the protein database generated by Kaiko, we re-analyzed the mass spectra from the soil samples using the database search tool MSGF+ and identified 30,762 unique peptides from 31,848 PSMs with 5% peptide FDR (see Methods). We performed functional annotations with these identified peptides using Unipept^29^, and found 1,760 Enzyme Commission (EC) numbers matched to 11,646 peptides (42%). Functions in the top 20 EC numbers (Supplementary Table 2) included various enzymatic functions for transcription and translation, energy production and signaling. 787 EC numbers were mapped to KEGG metabolic pathways, extensively covering carbohydrate and amino acid metabolism, as well as the metabolism of cofactors, vitamins and xenobiotics.

Among identified peptides, 14,028 peptides were highly conserved sequences and therefore were assigned to bacterial phyla. 3,228 of these phyla-affiliated peptides were linked to 708 EC numbers (Supplementary Figure 4). These highly conserved peptides were assigned to ubiquitous bacterial functions commonly detected across most phyla, such as DNA-directed RNA polymerase (EC:2.7.7.6) and H(+)-transporting two-sector ATPase (EC:7.1.2.2), With NAD(+) or NADP(+) as acceptor (EC:1.2.1.−), Acting on ATP (EC:3.6.4.-), Protein-synthesizing GTPase (EC:3.6.4.-). In particular, EC numbers of highly-ranked peptide counts were mainly detected in abundant phyla (Proteobacteria, Actinobacteria, and Acidobacteria) and functional information was biased by the common and abundant proteins.

We next examined the mapped EC numbers to identify metabolic functions for specific taxa (Fig. 3). By mapping the taxonomic affiliation of the enzymatic reactions within metabolic pathways it was possible to determine which metabolic pathways were shared or unique among the represented phyla. EC numbers involved in carbon metabolism were often found in organisms from multiple phyla, and represent basic functions from glycolysis, carbon fixation, the TCA cycle, etc. Enzymes and metabolic functions for 427 EC numbers were represented by only a single phylum, and are shown with different colors in Figure 3. The two most abundant phyla detected in the metaproteomics data were Proteobacteria and Actinobacteria. It is clear from the functional mapping of peptides that these two phyla utilize distinct metabolic routes. For example, purine metabolism contains numerous enzymes which are exclusively found in either Actinobacteria or Proteobacteria (Supplementary Fig. 5). Significant divergence between these two dominant taxa was also seen in enzymes related to amino acid metabolism. Unique metabolic capacity is also observed for organisms with less proteomic coverage. Despite having relatively few identified peptides, Verrucomicrobia were the only species with enzymes for folate metabolism.

**Figure 3.**
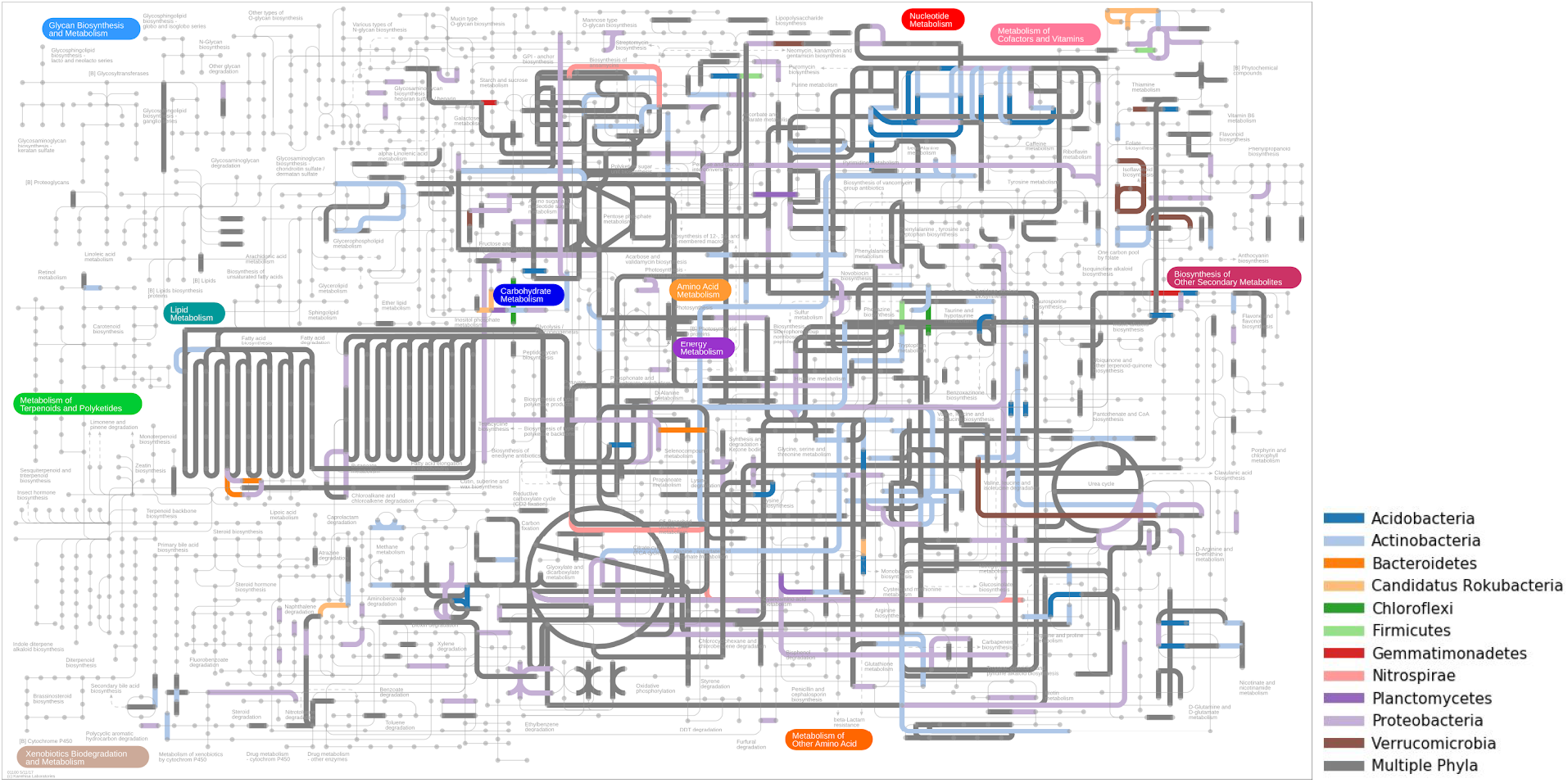
Distribution of bacterial functions in the metabolic pathway map. Several metabolic steps are shared among multiple phyla (dark grey). Other colors indicate unique EC numbers and their associated metabolic function found only in a specific phylum.

Finally, we examined the peptides and biological functions associated with species unique to the Kaiko database, i.e. species not found in the 16S rRNA sequences. 266 peptides were identified in Candidatus Rokubacteria and mapped to EC numbers. Biological functions associated with six EC numbers were exclusive to Candidatus Rokubacteria; 4-hydroxy-tetrahydrodipicolinate reductase (EC:1.17.1.8, lysine biosynthesis and monobactam biosynthesis), pyrroloquinoline-quinone synthase (EC:1.3.3.11), thioredoxin-disulfide reductase (EC:1.8.1.9, selenocompound metabolism), 3-oxoadipate enol-lactonase (EC:3.1.1.24, benzoate degradation), inositol-phosphate phosphatase (EC:3.1.3.25, inositol phosphate metabolism and streptomycin biosynthesis), and aminopyrimidine aminohydrolase (EC:3.5.99.2, thiamine metabolism).

## Discussion

Although genome and metagenome sequencing have greatly expanded the number of species that contain a sequenced genome and therefore an annotated proteome, there are still significant practical and financial barriers that prevent labs from always having an assembled and well-annotated genome for samples taken from nature. Yet metaproteome spectrum identification tools rely on a protein sequence database. Therefore tools which can create a proteome database for environmental samples without requiring sequencing data are a significant benefit to the microbiome community. One option for creating a proteome database without using sequencing data utilizes a *de novo* interpretation of metaproteomics data to identify organisms present in the sample. A significant drawback of current *de novo* tools is their poor performance on spectra from diverse organisms (see Figure 1). Algorithms which are only exposed to a limited number of organisms^11,12^, or those that focus only on human data^30^, will be inadequate when faced with the vast sequence diversity of microbial proteins found in soil and environmental samples.

To assist in the analysis of metaproteomic data, we have created a pipeline for generating the proteome sequence database directly from the metaproteomic data. A key element in our pipeline is a new *de novo* spectrum annotation tool, Kaiko, which has significantly improved accuracy compared to other *de novo* algorithms. This improvement comes from a deliberate focus on training the algorithm with mass spectrometry data from dozens of diverse environmental bacteria. Moreover, our training dataset size is dramatically larger than comparable *de novo* tools in terms of the number of peptides and spectra, which was essential for overcoming an overfit model. We evaluated Kaiko by using it to identify the taxonomy of bacterial soil isolates, including samples from phyla where no training data existed. Thus, it is better equipped for evaluating metaproteomics data where identifying spectra from diverse organisms is essential.

When using Kaiko as part of our database generation pipeline to identify soil community composition, we were able to identify all abundant species from 16S data, and also new species with significant proteomic evidence which were not seen in the sequencing data. Indeed, five of the top sixteen taxa (>30%) identified in the metaproteomics data were not identified in sequencing data. These ‘hidden microbes’ represent bacteria that are known to play an important role in community metabolism and function^23^, including secondary metabolite biosynthesis^31,32^ as was seen in our Candidatus Rokubacteria data. We also note that the metaproteomics pipeline was able to identify fungi in the soil, which are entirely absent in 16S data.

A second significant advantage of inferring community composition directly from metaproteome data is the level of taxon specificity. Using metaproteome data, we could narrow taxon identification to species or strain (98%). However taxa identified using 16S data for these same samples frequently were only able to distinguish broad taxonomic levels. Unfortunately, spectrum identification algorithms generally suffer a significant sensitivity loss when working with large protein databases^17^. Therefore, methods which specify community composition in broad taxonomic terms will yield poor results, compared to a method which is able to narrowly define organisms present in the community.

As metaproteomics data analysis continues to mature, progress will happen in multiple areas, e.g. more sensitive peptide ID algorithms, improved protein inference for multi-organism mapped peptides and functional analysis of pathways with multiple participating organisms. But a central feature in all of this work is the original identification of spectra, and currently the best algorithms require a protein database. Thus the creation of a protein sequence database is a pivotal step in metaproteomics data analysis. The most important future improvement in creating a protein sequence database will come from greater coverage and greater specificity in the identification of community membership. *De novo* proteomics offers one avenue for this, which is independent of advances made in sequencing technologies. Improving the accuracy of *de novo* tools, especially with regard to diverse environmental sequences, will be a significant benefit to metaproteomics.

## Acknowledgements

The authors thank Court Corley and Nathan Hodas (PNNL) for insightful discussions. We thank Kristen DeAngelis and Grace Pold (University of Massachusetts Amherst) for natural isolate samples. Funding for this project was provided by the U.S. Department of Energy, Office of Science, Office of Biological and Environmental Research, Early Career Research Program (to SHP and KEBJ) and PNNL’s Deep Learning for Scientific Discovery initiative. Proteomics data used in this manuscript were generated in the Environmental Molecular Science Laboratory, a DOE national scientific user facility at Pacific Northwest National Laboratory (PNNL) in Richland, WA. Battelle operates PNNL for the DOE under contract DE-AC05-76RLO01830.

## Methods

### Data generation for Kaiko

#### Cell culture and sample preparation

The growth, sample preparation and data collection was reported previously^33^. Cells were harvested by centrifuging at 3,500 × *g* for 5 min at room temperature and washed twice with 5 mL PBS by centrifuging at the same conditions. Cells were lysed in a Bullet Blender (Next Advance) for 4 minutes at speed 8 in 200 μL of 100 mM ammonium bicarbonate (NH_4_HCO_3_) and approximately 100 μL 0.1 mm zirconia/silica beads at 4° C. Lysates were transferred into clean tubes and the remaining beads were washed with 200 μL of 100 mM NH_4_HCO_3_. The supernatants from the washing step were collected and combined with the cell lysate. Resulting protein extract was assayed by bicinchoninic acid (BCA) assay (Thermo Fisher Scientific, San Jose, CA) following manufacturer instructions. Aliquots of 300 μg of proteins were denatured and reduced using 8M urea and 5 mM DTT, and incubated at 60° C for 30 min with 850 rpm shaking. Samples were then diluted 10 fold in 100 mM NH_4_HCO_3_ and CaCl_2_ was added to a final concentration of 1 mM using a 1M stock. Trypsin was added at 1/50 of the protein concentration and the digestion was carried out for 3 h at 37° C. Digestion products were desalted in 1-mL C18 cartridges (50 mg beads, Strata, Phenomenex). Cartridges were activated with 3 mL of methanol and equilibrated with 2 mL of 0.1% TFA before loading the samples. After sample loading, the cartridges were washed with 4 mL of 5% acetonitrile (ACN)/0.1% TFA and peptides were eluted with 1 mL of 80% ACN/0.1% TFA. Peptides were dried in a vacuum centrifuge, resuspended in water and assayed using a BCA assay. Peptide concentrations were normalized to 0.1 μg/μL before randomization and analysis by liquid chromatography-tandem mass spectrometry (LC-MS/MS).

#### LC-MS/MS data acquisition

The data acquisition was performed as previously described in detail^33^ using a Waters nanoEquity™ UPLC system (Millford, MA) coupled with a Q Exactive Plus mass spectrometer from Thermo Fisher Scientific (San Jose, CA). The LC was configured to load the sample first on a solid phase extraction (SPE) column followed by separation on an analytical column. 500 ng of peptides were loaded into the SPE column (5 cm × 360 μm OD × 150 μm ID fused silica capillary tubing (Polymicro, Phoenix, AZ); packed with 3.6-μm Aeries C18 particles (Phenomenex, Torrance, CA) and the separation was carried out in a capillary column (70 cm × 360 μm OD × 75 μm ID packed with 3-μm Jupiter C18 stationary phase particles (Phenomenex). The elution was performed at 300 nl/min flow rate and the following gradient of acetonitrile (ACN) in water, both containing 0.1% formic acid: 1-8% ACN solvent in 2 min, 8-12% ACN in 18 min, 12-30% ACN in 55 min, 30-45% ACN in 22 min, 45-95% ACN in 3 min, hold for 5 min in 95% ACN and 99-1% ACN in 10 min. Eluting peptides were directly analyzed in the mass spectrometer by electrospray using etched silica fused tips^34^. Full MS spectra were acquired at a scan range of 400-2000 m/z and a resolution of 35,000 at m/z 400. Tandem mass spectra were collected for the top 12 most intense ions with ≥ 2 charges using high-collision energy (HCD) fragmentation from collision with N2 at a normalized collision energy of 30% and a resolution of 17,500 at m/z 400. Each parent ion was targeted once for fragmentation and then dynamically excluded for 30 s.

#### Peptide identification for training/testing the Kaiko model

In the training and test set, the true source\taxonomy of each sample is known. To create the ground truth of spectrum identifications, we used the correct organism’s protein sequence database and annotated spectra with the MSGF+ algorithm, as previously described^33^. PSM results from MSGF+ were filtered using a q-value threshold of 0.001. The PSMs passing this filter were considered the ground truth for the deep neural network training and testing. Because our use of this data is for *de novo* spectrum annotation, we limited peptides/spectrum matches further to exclude peptides longer than 30 residues as these were unlikely to have complete peptide fragment peaks, which are important for a *de novo* solution. We also filtered peptides with a precursor mass >3000 Da. After filtering, the total number of distinct peptides was 1,013,498 from 5,116,305 spectra. Peptide sequences are highly specific to each organism, and the overlap between organisms was very low. Except for the pairs of organisms within the same genus or species (i.e. the two different strains of *B. subtilis* or the two different species within *Bifidobacterium*), the average amount of shared peptides between any two organisms was ~0.17%. These arise from highly conserved proteins like EF-Tu or RpoC for which peptides can be found conserved across phyla.

### Training Kaiko

#### Codebase

Kaiko is based on DeepNovo, a deep neural network algorithm for peptide/spectrum matching^12^. We downloaded the source code for DeepNovo (https://github.com/nh2tran/DeepNovo) and its pre-trained model, which is publicly available at https://drive.google.com/open?id=0By9IxqHK5MdWalJLSGliWW1RY2c. As described below, we modified the original DeepNovo codebase, keeping with Python 2.7 and TensorFlow 1.2 as used in the original. First, we modified the codebase to accept multiple input files for training and testing. Our training and testing data came from over 250 mass spectrometry files, but the original DeepNovo was designed for only a single input file. Therefore, we added extra command-line options (e.g., --multi_decode and --multi_train) and the associated wrapper methods to allow for multi-file execution. A second change was done to avoid rebuilding the Cython codes on every parameter adjustment. For this, we replaced the Cython with the python *numba* package without any loss of performance and speed. Finally, we changed the code for spectral modeling based on domain knowledge. Specifically, we corrected the mass calculation of doubly charged ions and changed the bins used for isotopic profiles within the ion-CNN model.

We trained multiple models for Kaiko, which differed primarily in the number of peptides/spectra used during training: ~300K spectra, 1M spectra, 2M spectra, 3M spectra and the final models trained with all spectra. When training the final model on the full dataset, we adjusted the learning rate to 10^−4^ rather than using the default value (10^−3^) of AdamOptimizer in DeepNovo. Training our final model requires very significant computational resources and time. With the hardware used in this project, training took ~12 hours per epoch; our final model was achieved after 60 epochs. All training and testing was performed on PNNL’s Marianas cluster, a machine learning platform that is part of PNNL’s Institutional Computing. System specifications on the nodes used in this training were: Dual Intel Broadwell E5-2620 v4 @ 2.10GHz CPUs (16 cores per node), 64 GB 2133Mhz DDR4 memory, and Dual NVIDIA P100 12GB PCI-e based GPUs.

#### Experimental Design and Statistical Rationale

Given that Deep Neural Networks are very sensitive to overfit during the training procedure, we anticipated that a very large amount of data would be required to make useful models. The original DeepNovo was trained on 50,000 spectra and we believed that a significantly larger amount of data would be necessary. As described below in “Assessing Progress” we were able to quickly determine that a model with only 300,000 spectra was overfit. We therefore determined that we would aim for 5,000,000 spectra representing about 1,000,000 peptides in order to have sufficient data for training the very large neural networks that comprise Kaiko. During training we were able to determine that this number was more than sufficient to produce a generalized model that did not overfit to training data. Spectra included in the training, validation and testing set are assessed as described above in the “Data Generation” section.

#### Assessing Progress

The training regimen for deep learning is pragmatically broken up into several rounds of iteration over the training data, called epochs. During each epoch, a mini-batch stochastic optimization was employed, in which each batch of 128 spectra is randomly chosen and training proceeds on each batch one at a time. The model is trained by updating the parameters within the neural network (weights and biases) after each batch is compared to the true labels. While training, the error associated with the model can be calculated as a cross-entropy loss for the probabilities of correctly predicting the amino acid letters on the training data. After each batch, we also randomly sample 15,000 spectra from the validation dataset (~1% of total testing data) and compute the loss error, which we call the validation error. Importantly, model performance on this validation set is **not used** to update the model parameters; we simply use it to independently evaluate model performance and make a checkpoint to track the best models. The training and validation error after each batch for 20 epochs of training is shown in Supplementary Fig.2.

By comparing the training and validation error, we clearly see when the model has started to overfit. This happens when the training error crosses over (becomes smaller than) the validation error and continues to decrease as the validation error levels off. This is a result of the model learning specific features of the training data that are not generalizable. In models built with more than 3 million spectra, no overfitting is seen yet; models built with less than 3 million spectra quickly overfit to the training data.

### Comparing Kaiko to other *de novo* tools

To compare the performance of Kaiko to state-of-the-art *de novo* tools, we analyzed all files in the testing data sets using DeepNovo^12^, PEAKS^35^ and Novor^30^. As mentioned above, we used a pre-trained model for the DeepNovo to predict peptide sequences for the test files using a ‘decode’ option. PEAKS Studio version 8.5 was run using default data refinement options on mzML formatted data^36^. *De novo* settings were as follows: precursor error tolerance - 20ppm, fragment ion error tolerance - 0.02 Da. Oxidation of methionine was set as a variable modification. For Novor the spectral files were converted from mzML to Mascot generic format (MGF) using MSConvert^37^. Novor version 1.05 was run using the following settings: fragmentation - HCD, massAnalyzer - FT, precursor error tolerance - 20ppm, fragment ion error tolerance - 0.02 Da. Oxidation of methionine was set as a variable modification. All other settings were left at their defaults. Only the best peptide spectrum match was used in the evaluation. Please refer to https://github.com/PNNL-Comp-Mass-Spec/Kaiko_Publication/analysis/for_novor and /for_peaks for specific implementation details.

### Assigning taxonomy to unknown samples

Proteomics data from six bacterial soil isolates was acquired using the same sample preparation and LC-MS/MS method as described above. The isolates are from the natural isolate collection at the Kristen DeAngelis laboratory at the University of Massachusetts Amherst, and researchers at PNNL were blinded to the identity of these isolates until after both data generation and analysis were finished. Kaiko’s top-scoring peptide sequence for each spectrum was used for species identification. We filtered these peptide/spectrum matches to include only the top 25% according to Kaiko’s quality prediction score. We then exclude sequences shorter than 10 and longer than 17 residues. The resulting sequences were used to search the Uniref100 protein database [https://www.uniprot.org/uniref/] using DIAMOND^13^ to identify an organism(s) containing that peptide sequence. Only database matches of 100% were retained for species prediction. Taxon scoring then proceeded using a two-pass procedure. In the first pass, for each peptide sequence, all taxa possessing a 100% match were assigned 1 hit, such that multiple taxa were often assigned a hit from a single peptide sequence. Taxa were then ranked by the total number of hits assigned. In the second pass, hits were only assigned to the highest-ranking taxon with a 100% match to each predicted sequence. In this way, scoring is assigned to the candidate most likely to be correct.

### Metaproteomics data analysis

#### Sample preparation from soils

Kansas prairie soil was quickly thawed and weighed into 10 g aliquots in 50 mL methanol/chloroform compatible tubes (Genesee Scientific, San Diego, CA) along with 10 mL of 0.9–2.0mm stainless steel beads, 0.1mm zirconia beads and 0.1mm garnet beads. All beads had previously been washed with chloroform and methanol and dried in a fume hood. Protein extraction occurred using a modified method of the Folch extraction^38^ specifically for soil called Soil MPLex (Metabolite, protein, lipid extraction)^39^. Here, 4 mL of ice-cold ultrapure “Type 1” water (Millipore, Billerica, MA) was added to each sample and transferred to an ice bucket in a fume hood. Using a 25 mL glass serological pipette, −20 °C 2:1 chloroform: methanol (v/v) (Sigma-Aldrich, St. Louis, MO), was added to the sample in a 5:1 ratio over sample volume (20 mL) and vigorously mixed (by vortexing). The tubes were attached to a 50 mL tube vortex-attachment and horizontally mixed for 10 min at 4 °C and placed inside a −80 °C freezer for 5 min. Using a probe sonicator (model FB505, Thermo Fisher Scientific, Waltham, MA) inside a fume hood, each sample was sonicated with a 6mm probe (20 kHz fixed ultrasonic frequency) at 60% of the maximum amplitude for 30 s on ice, allowed to cool on ice, then sonicated once more. Samples were allowed to cool for 5 min. at −80 °C, then mixed for 60 s and centrifuged at 4,500 *xg* for 10 min at 4 °C. The upper aqueous phase was removed and the interphase containing proteins that partitioned between the methanol and chloroform phases was collected into a separate tube and precipitated through addition of 5 mL of −20 °C 100% methanol. Following methanol addition, the tube was mixed then centrifuged at 4,500 *xg* for 5 min at 4 °C in order to pellet the proteins. The supernatant was decanted and the protein pellet dried upside down. Meanwhile, the bottom organic phase was removed, and 5 mL of −20 °C 100% methanol was added to the bottom debris pellet, mixed and centrifuged at 4,500 *xg* for 5 min at 4 °C. The supernatant was removed, and the protein pellet was dried upside down.

Protein pellets from both the debris and interphase were frozen and lyophilized for 2 h. Proteins from the interphase were solubilized by addition of 10 mL of SDS-Tris buffer containing 4% sodium dodecyl sulfate (SDS), 100mM DL-dithiothreitol (DTT) in 100mM Tris-HCl, pH 8.0, (Sigma-Aldrich, St Louis, MO), briefly probe sonicated at 20% amplitude, then incubated on a lab tube rotator for 30 min at 300 rpm, 50 °C. Proteins from the debris pellet were solubilized in 20 mL of SDS buffer, horizontally vortexed for 10 min. to lyse any remaining intact cells, then combined with the interphase proteins and mixed on the rotator assembly for the time remaining (approximately 20 min). Following mixing, the tubes were centrifuged at 4,500 *xg* for 10 min., and the supernatant from each tube were combined into a single 50 mL tube. The proteins were precipitated by adding up to 25% trichloroacetic acid (TCA; Sigma-Aldrich, St. Louis, MO), mixed and placed at −20 °C overnight. The proteins were thawed and centrifuged at 4,500 *xg* at 4 °C for 10 min to collect the precipitated proteins. The supernatant was gently decanted, and the protein pellet washed through addition of 2 mL of −20 °C acetone, mixed, then placed at −80 °C for 5 min. Proteins were pelleted by centrifugation for 10 min at 4,500 *xg* at 4 °C. The acetone was removed by gently decanting, and the wash step was repeated 2 more times. The washed pellet was then air dried by inverting the tube. After drying, 100 μl–200 μl of SDS-Tris buffer was added and the solution was transferred into 1.5 mL tubes and incubated at 95 °C for 5 min, then cooled at 4 °C for 10 min. The samples were centrifuged at 15,000 *xg* for 10 min to pellet any remaining debris and transferred into fresh 1.5 mL tubes in preparation for digestion using the Filter-Aided-Sample-Preparation (FASP) digestion method^40^. For protein digestion, up to 30 μl of proteins in SDS-Tris buffer were transferred to a 30,000 Da molecular weight cut off (MWCO) 500 μl spin filter provided in the Expedeon FASP kit (Expedeon LTD, Cambridgeshire, UK) along with 400 μl of 8 M urea solution. The spin filter was centrifuged at 14,000 *xg* for 30 min. The waste was removed from the collection tubes and 400 μl of 8M urea solution was added to each sample and centrifuged as described above, then repeated for a total of 3 urea additions. 400 μl of 25mM NH_4_HCO_3_, pH 8, was added and centrifuged as described above, then repeated for a total of 2 ammonium bicarbonate washes. The spin column was transferred into a fresh-labeled collection tube and 75 μl of NH_4_HCO_3_ was added to the filter along with 4 μl of 1 μg/μl molecular grade trypsin (Thermo Fisher, Waltham, MA) then incubated at 37 °C for 3 h. After digestion, 40 μl of NH_4_HCO_3_ was added to the sample and centrifuged at 14,000 *xg* for 20 min. Another 40 μl of NH_4_HCO_3_ was added to the top of the filter, mixed and centrifuged again for 10 min. The filter was discarded, and the collected peptides were treated with potassium chloride (KCl) in order to ensure all the SDS was removed^41^. To accomplish this, potassium chloride was added to the peptides in NH_4_HCO_3_ resulting in a final concentration of 2M KCl, then mixed and allowed to rest for 10 min. at room temperature. To pellet the SDS, the peptide solution containing NH_4_HCO_3_ and KCl was centrifuged at 14,000 *xg* for 10 min. The supernatant was transferred to a fresh tube without disturbing the SDS pellet and salts removed using a microspin C18 column according to the manufacturer’s instructions (the Nest Group, Inc., Southborough, MA). Peptides from the aliquots of 10 g of soil were combined to generate a single peptide sample. A bicinchoninic acid (BCA) assay (Thermo Fisher Scientific, Waltham, MA) was performed to determine the peptide concentration.

The peptide sample was separated with a commercial Waters (Milford, MA) XBridge 5 μm particle size C18 column, (4.6mm i. d. × 250mm length) with an attached 20mm long × 4.6mm i. d. guard column. Fractionation was performed at 0.5 mL/min using an Agilent 1100 series HPLC system (Agilent Technologies, Santa Clara, CA) with two mobile phases: A) 10mM NH_4_HCO_2_ (pH 10.0), and B) 10mM NH_4_HCO_2_ (pH 10.0) with acetonitrile (10:90). A six step gradient was adjusted over 120 min by replacing mobile phase A with B according to: 1) 100%–95% over the first 10 min., 2) 95%–65% from minutes 10 to 70, 3) 65%–30% from minutes 70 to 85, 4) then maintained mobile phase A at 30% from minutes 85 to 95, 5) re-equilibrating with 100% mobile phase A from minute 95 to 105, and 6) holding mobile phase A at 100% until minute 120. Fractions were collected every 1.25 min (96 fractions over the entire gradient) with every 24th fraction combined for a total of 24 final fractions (rows of a 96 well plate were pooled by every other row). All fractions were dried under vacuum and suspended in 25 μl H_2_O. A final BCA assay was done on the fractions and each were diluted to 0.1 μg/μl for LC-MS/MS analysis. (see LC-MS/MS data acquisition methods above).

#### Analyzing 16S rRNA amplicon sequences

16S rRNA gene amplicon sequencing data was downloaded from https://osf.io/4uvj7/, performed using the protocol developed by the Earth Microbiome Project^1^. Please refer to the previous studies^19,42^ for 16S rRNA gene amplicon sequencing in detail. The 16S rRNA amplicon sequences were first re-processed by Hundo pipeline^43^ (v1.2.8), a command line interface work comprising a set of existing software together with validated custom methods derived from QIIME^44^. In brief, the sequences were first quality filtered to remove the adaptors and contaminated reads from Phix genomes by BBDuk2^45^. The passing reads were merged and checked for chimera, which were subjected to be clustered into OTUs by VSEARCH^46^ using the default parameters. The abundance of each OTU was estimated by the read coverage of the OTU representative sequences (VSEARCH). In comparison to the Silva database^47^ implemented in Hundo, NCBI database was reported with a higher confidence of lineage assignment to lower taxonomy levels^48^. The de-replicated representative sequences of each OTU were then annotated following the same workflow coded in Hundo with modifications and using NCBI 16S Refseq database (https://www.ncbi.nlm.nih.gov/refseq/targetedloci/16S_process/, accessed on Apr 9th, 2020) instead. The top 25 hits of each OTU representative sequence were kept and screened for ones with percent identity higher than 85% and bit score greater than 125. For OTUs with more than one qualified hits, we will perform Lowest common ancestor (LCA) algorism using a R package, taxize^49^. OTUs with only one qualified hit adopted the lineage of the hits and the rest were left unclassified.

#### Constructing protein database for metaproteomic data analysis with Kaiko

Raw mass spectrometry files were converted to the PSI open format mzML^36^ using msConvert^37^, which were converted to MGF files compatible with the Kaiko model. After performing Kaiko prediction, as used for assigning taxa to the unknown samples, we used Kaiko’s top 25% scoring peptide sequences predicted from each sample to identify the most likely candidate organisms using DIAMOND over the Uniref100 database. The protein database was constructed by aggregating all the reference sequences associated with top 100 bacterial organisms from the Uniref100 into a single fasta (8.2GB).

#### Peptide identification and functional analysis with the constructed database

Against the protein database constructed from the Kaiko prediction, MSGF+ was performed to identify peptide sequences with the false discovery rate (FDR) cutoff. The search parameters and values or settings were as follows: PrecursorMassTolerance, 20.0 ppm; IsotopeErrorRange, −1,1; TargetDecoyAnalysis, true; FragmentationMethod, as written in the spectrum; InstrumentID, 0; Enzyme, Tryp; NumTolerableTermini, 2; MinPeptideLength, 6; MaxPeptideLength, 50; MinCharge, 2; MaxCharge, 5; and NumMatchesPerSpec, 1. PSM results from MSGF+ were filtered using MSnID (v1.20.0)^50^. Filters based on the cleavage patterns for the trypsin were applied, e.g., nuIrregCleavages==0 and numMissCleavages<=2. Optimizing the MS/MS filter was applied to achieve the maximum number of identifications within a given FDR upper limit threshold. Nelder-Mead method was employed for parameter optimization (MS-GF:SpecEValue and absParentMassErrorPPM), and for 5% peptide FDR, SpecEValue≤1.0e-11, and 11 ppm mass window with the ppm offset adjustment were determined. For functional annotation for metaproteomics, Unipept^29^ (v4.3.5, https://unipept.ugent.be/datasets, accessed on Jun 2nd, 2020) was used with “Equate I and L” and “Filter duplicate peptides” options.

### Data Availability

The mass spectrometry proteomics data for this benchmark set are split into two separate depositions, for the training and testing datasets respectively. The training dataset consists of spectra from 51 organisms and has been deposited to the ProteomeXchange Consortium via the PRIDE^25^ partner repository with the dataset identifier PXD010000. The testing dataset consists of spectra for 4 organisms and has been deposited to the ProteomeXchange Consortium via the PRIDE partner repository with the dataset identifier PXD010613. The metaproteomics dataset has been deposited to the MassIVE Repository with the accession identifier MSV000086336.

## Supplementary Figures

**Supplementary Figure 1.**
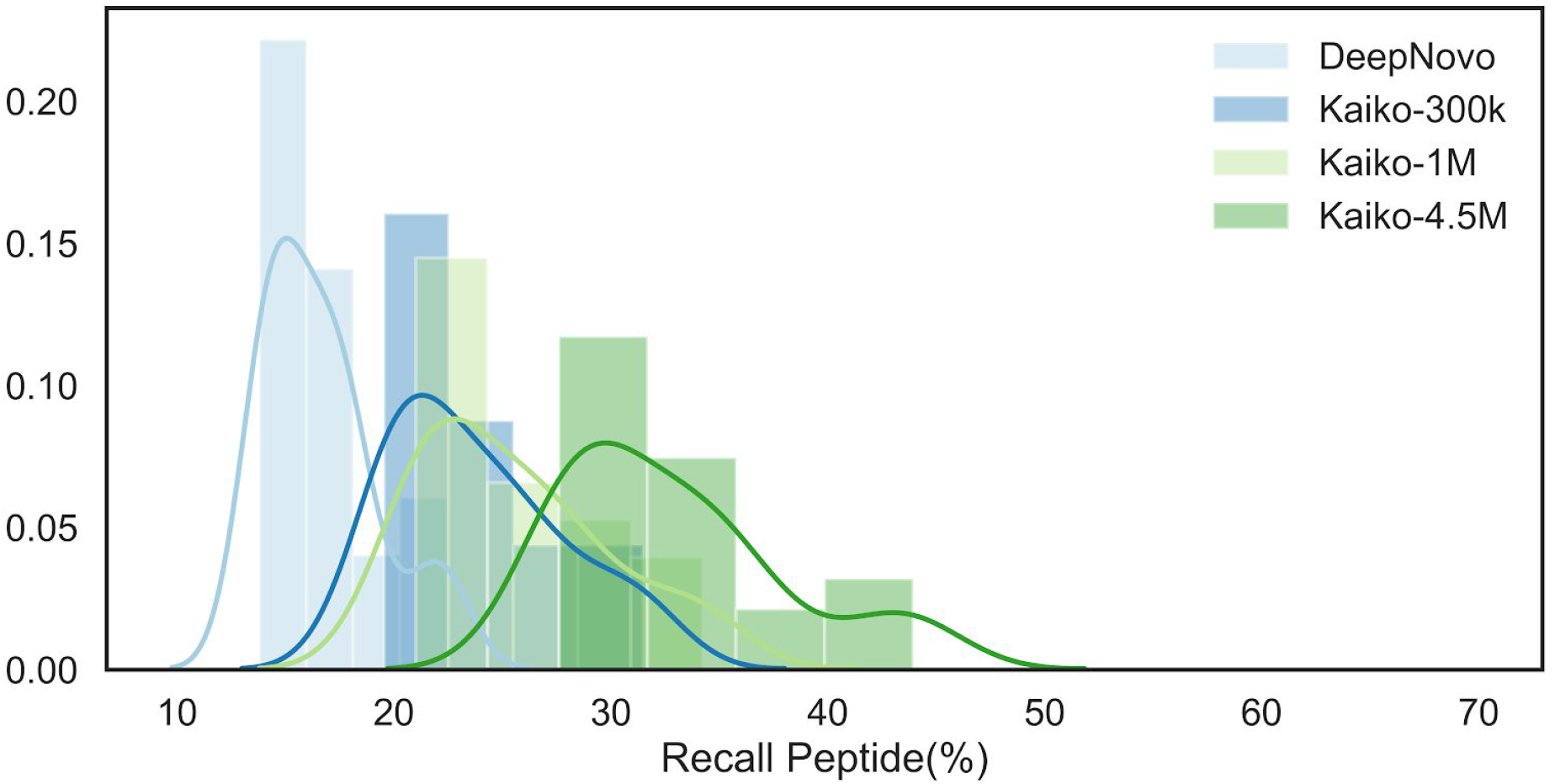
Improving deep neural networks with more training data. The accuracy of peptide/spectrum matching is shown for four deep neural network models. DeepNovo is a pre-trained publicly available model trained on 50,000 spectra. Kaiko was trained with varying numbers of spectra. The final model was trained with 4.5 million spectra. A significant improvement is seen in model performance with increased training data.

**Supplementary Figure 2.**
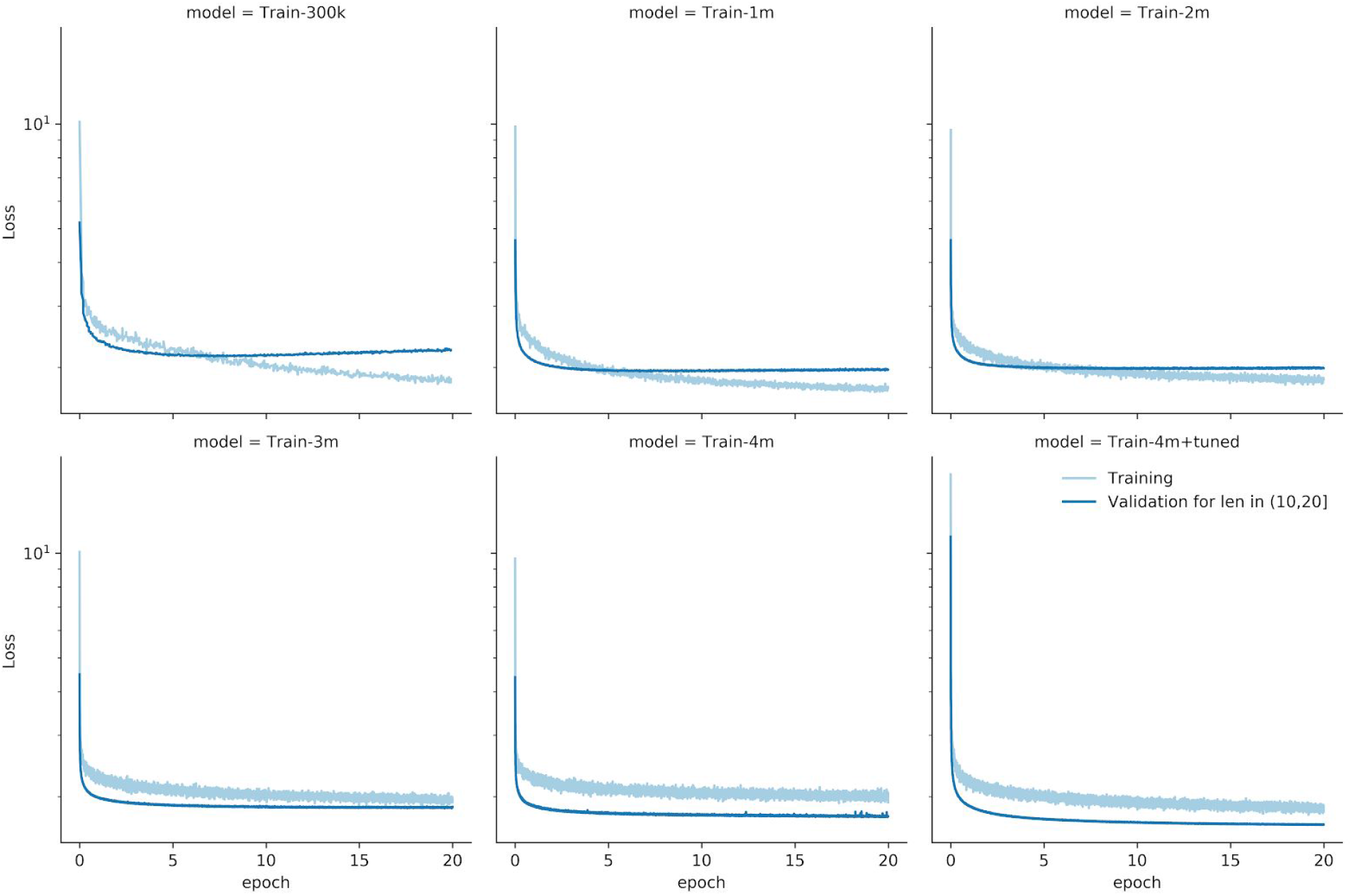
Training and validation error. During the epochs of learning for the deep neural network, progress is measured by evaluating the accuracy of spectrum annotation. We employ a cross-entropy loss function, which represents how well the algorithm is being trained, with small numbers being better. The light blue line represents the error on batches of training data. The dark blue lines represent the error on the random samples of the validation data. When the training error improves beyond the validation error, the model is likely to be overfitting.

**Supplementary Figure 3.**
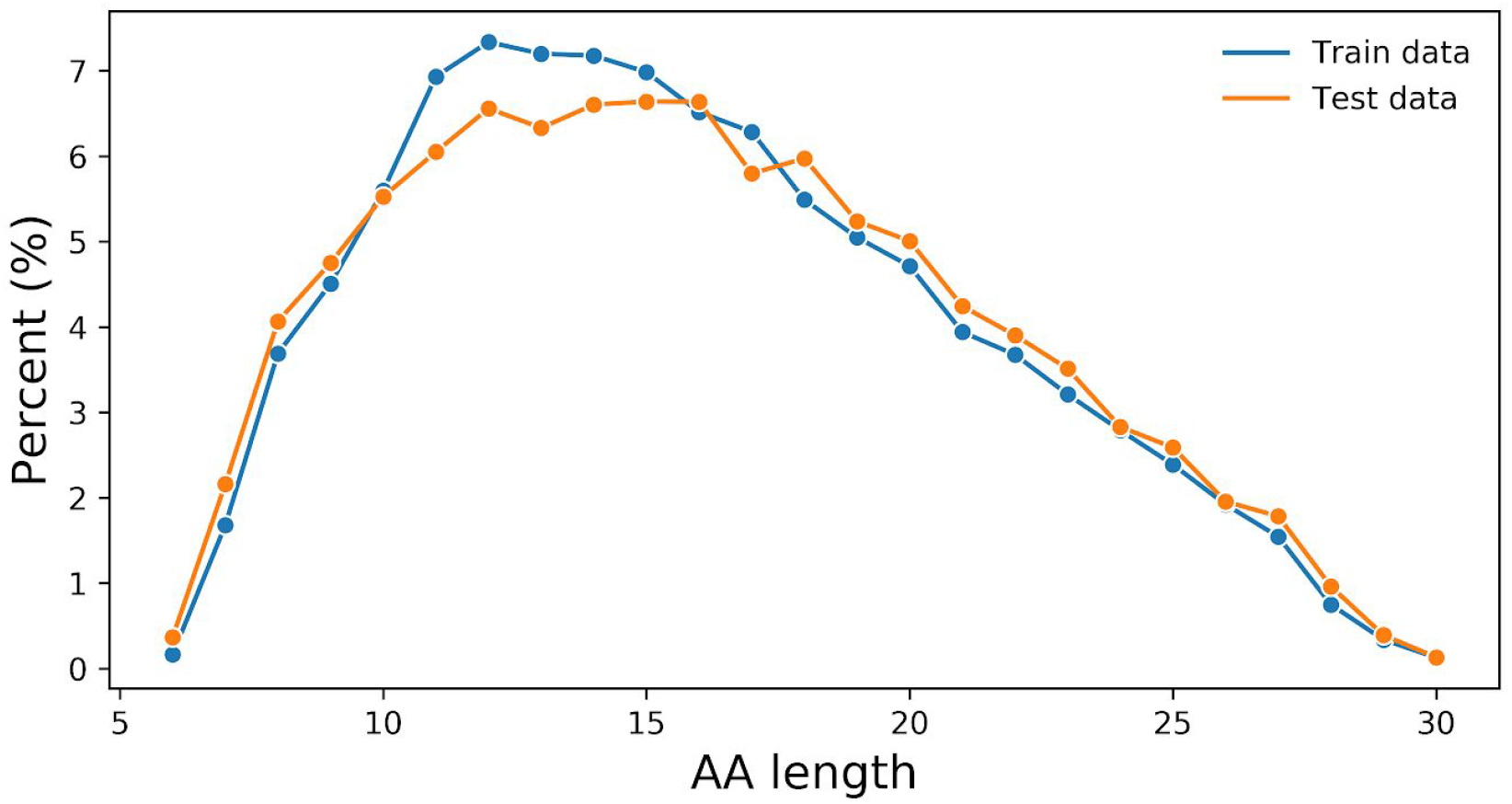
Distribution of peptide lengths. used for training and testing the Kaiko model.

**Supplementary Figure 4.**
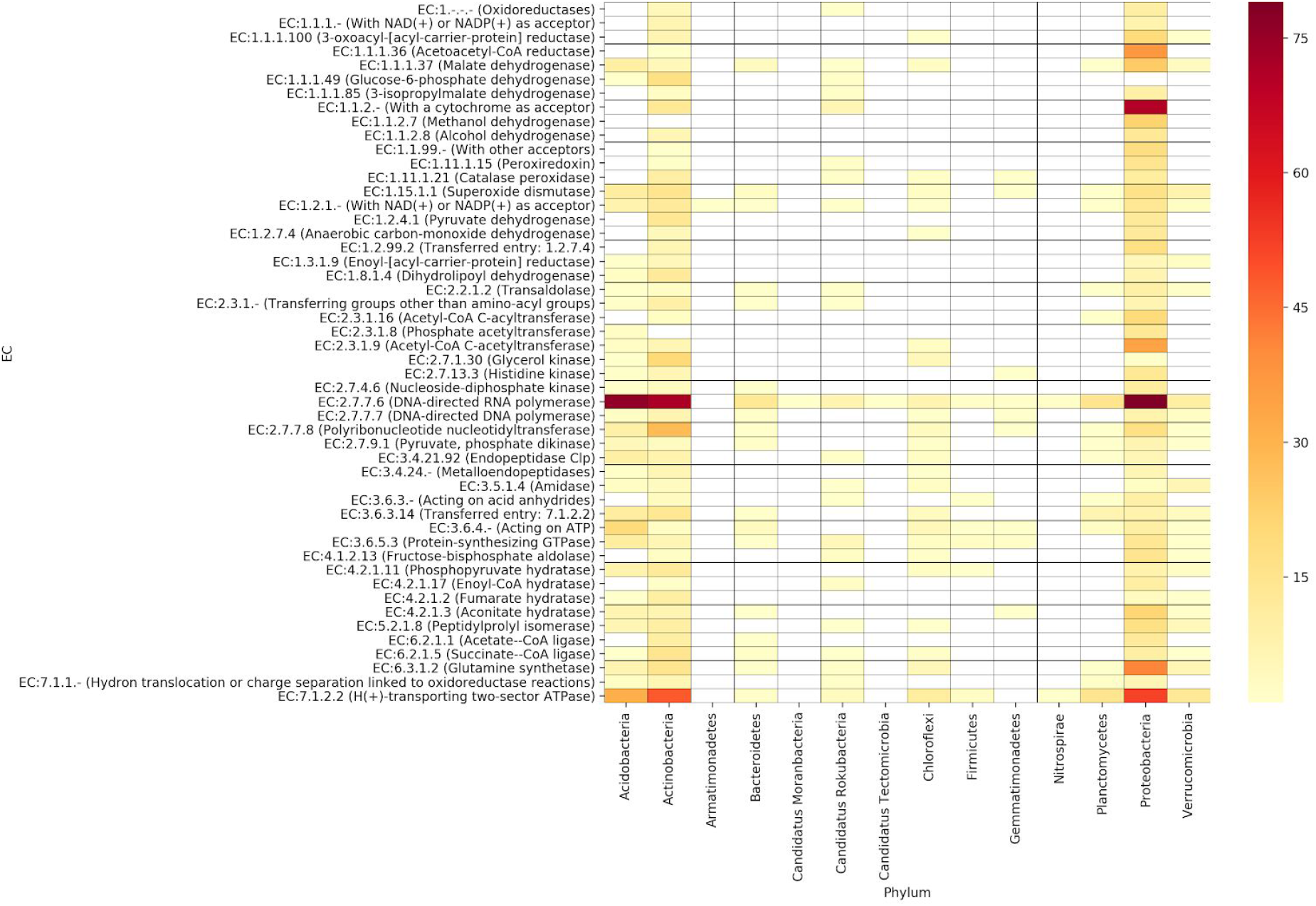
Heatmap of the peptide counts for the most common functions over the diverse phyla. Columns and rows in the heatmap represent the phyla and EC numbers, respectively. Cell colors indicate the number of phyla-affiliated peptides corresponding to a specific phylum and function.

**Supplementary Figure 5.**
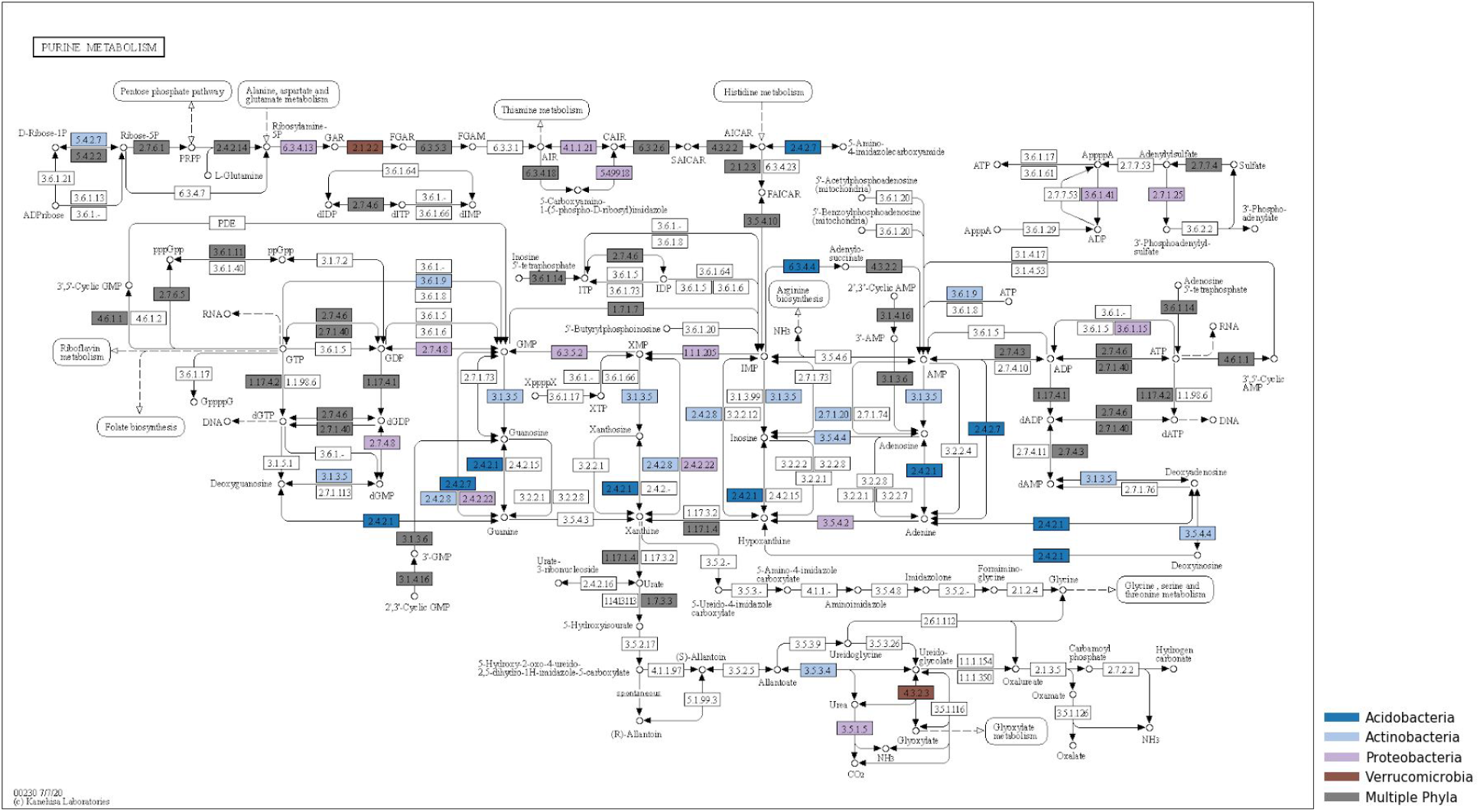
Taxa-specific peptides for enzymes in purine metabolism at the phylum level. Many EC numbers were uniquely detected in four phyla (Acidobacteria, Actinobacteria, Proteobacteria, and Verrucomicrobia).

## Supplementary Tables

**Supplementary Table 1.**
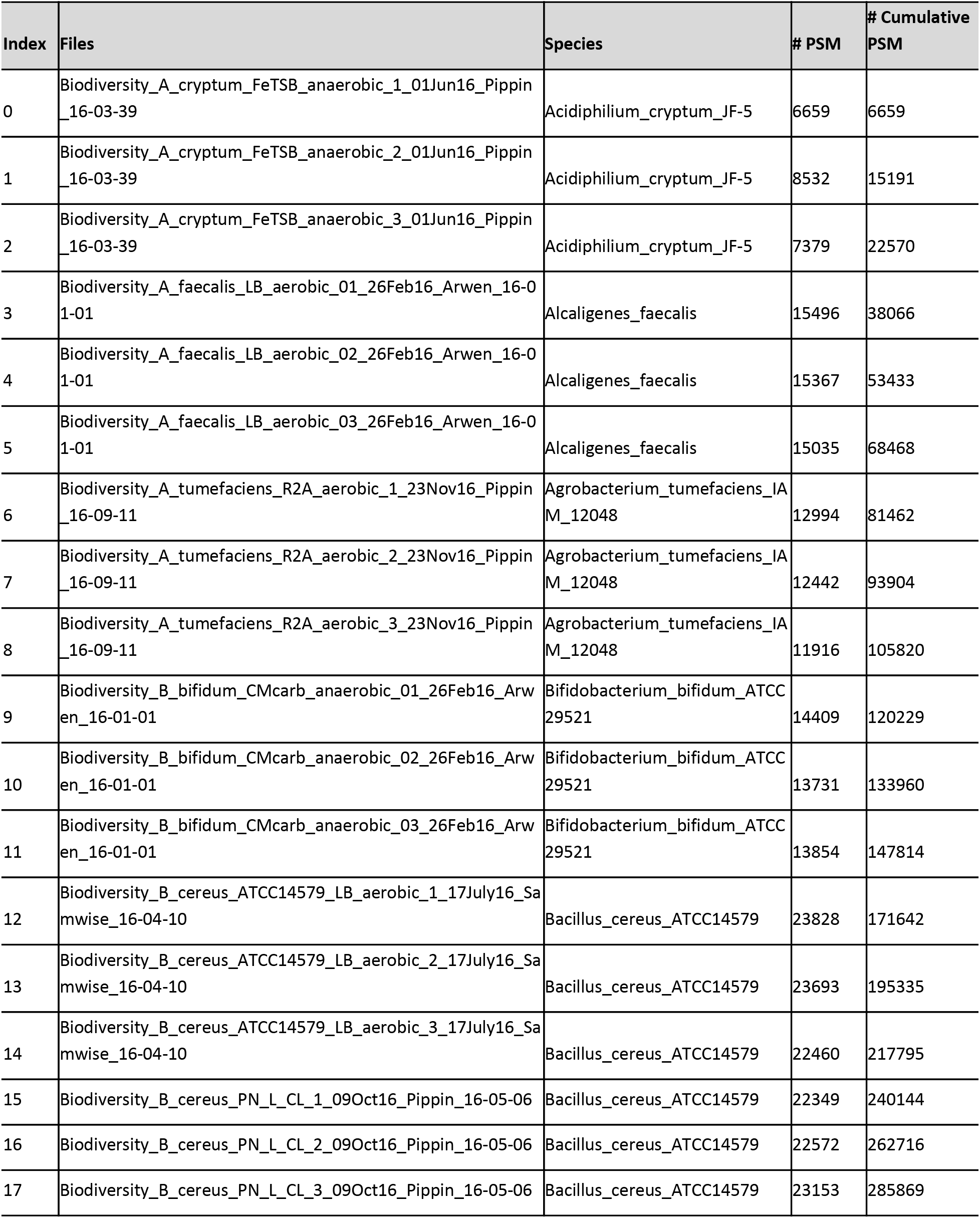
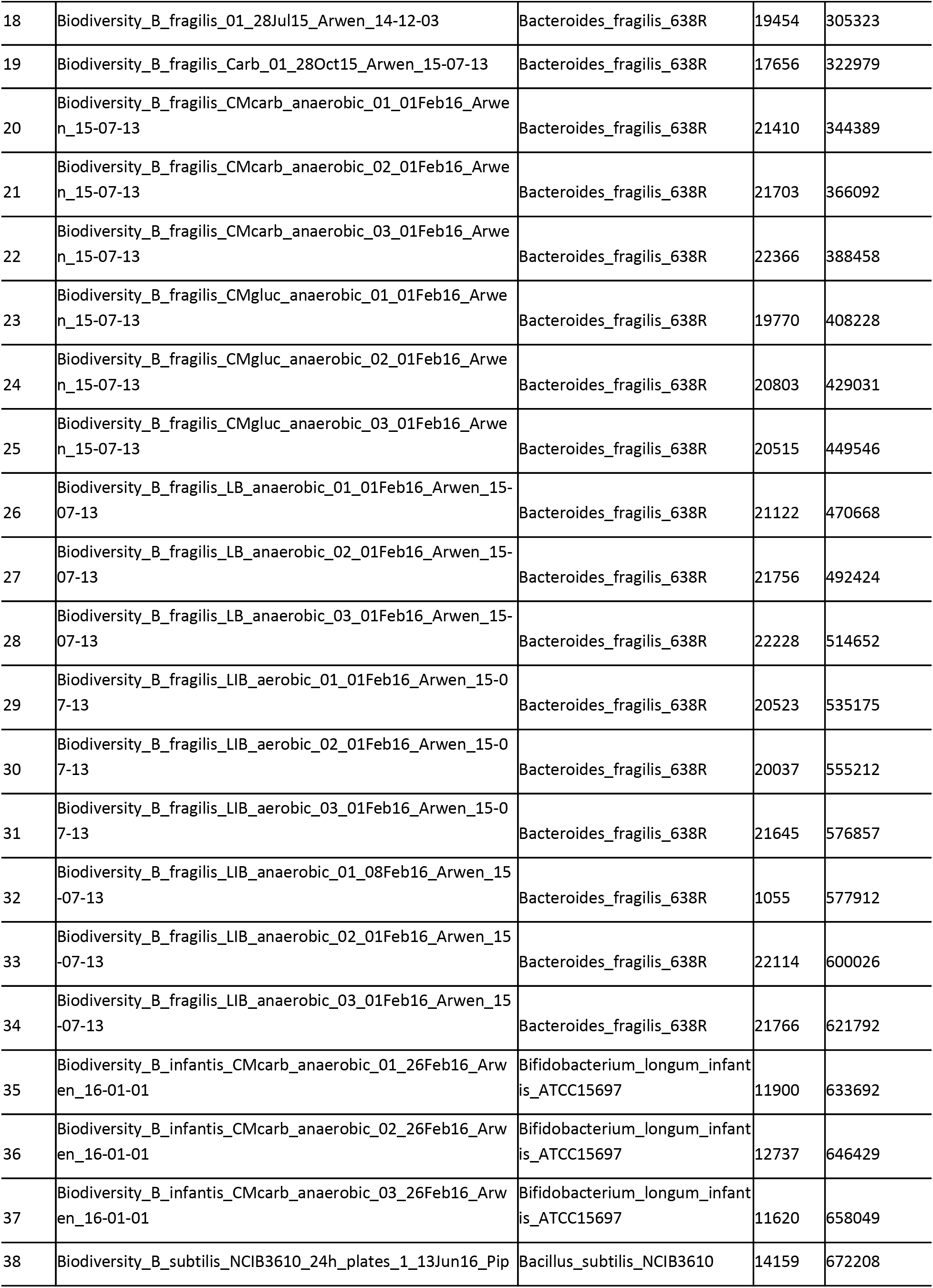
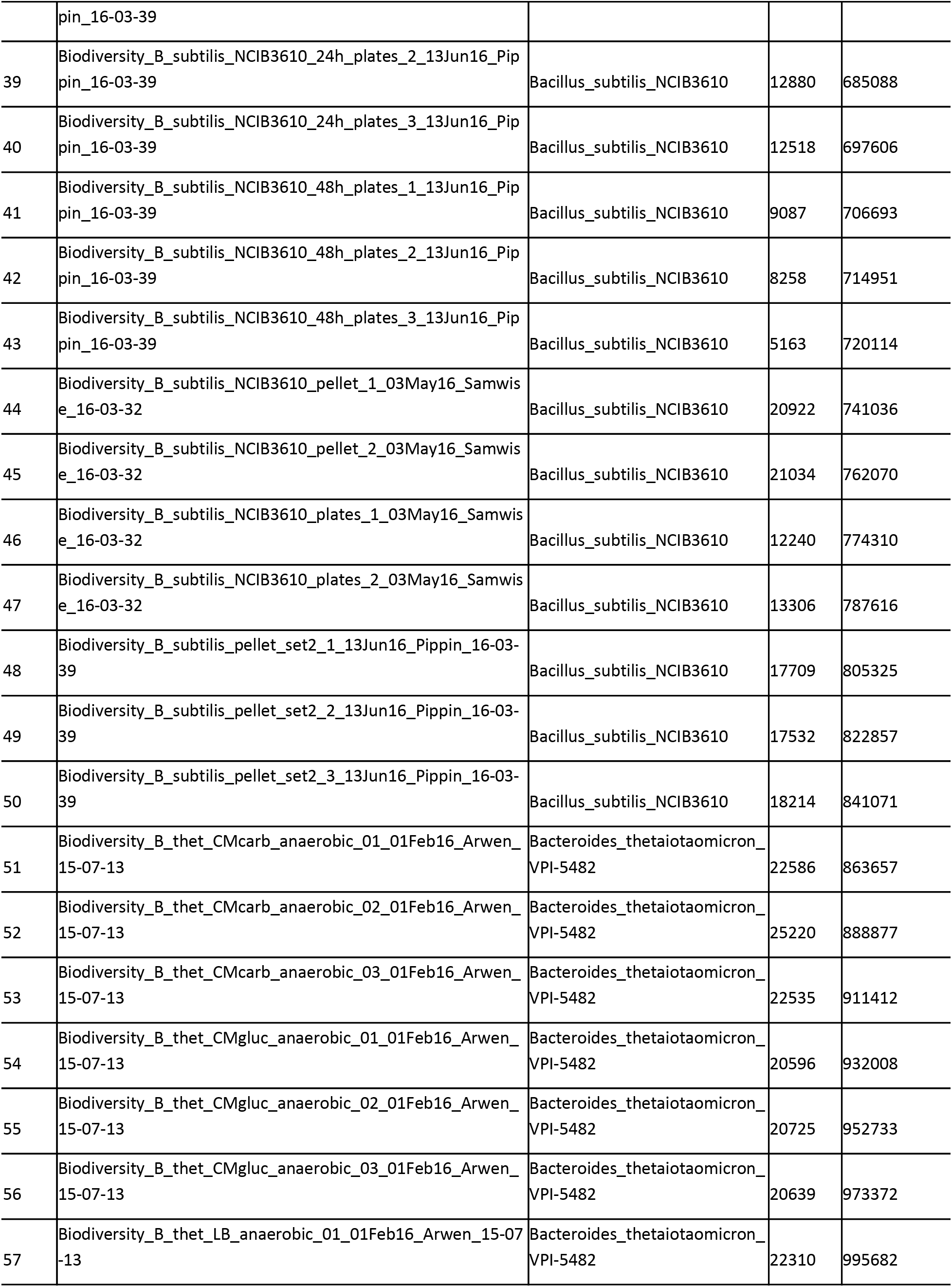
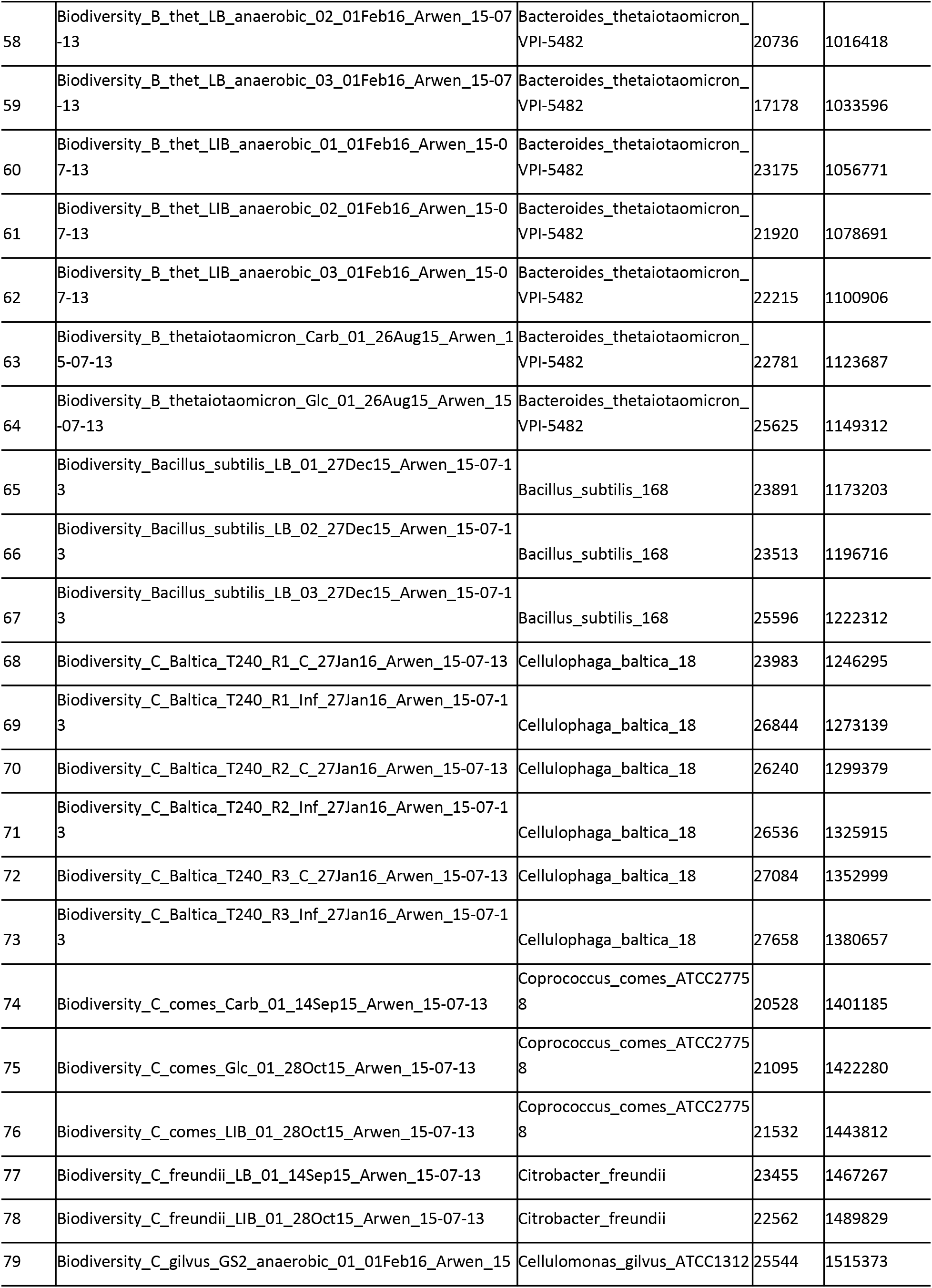
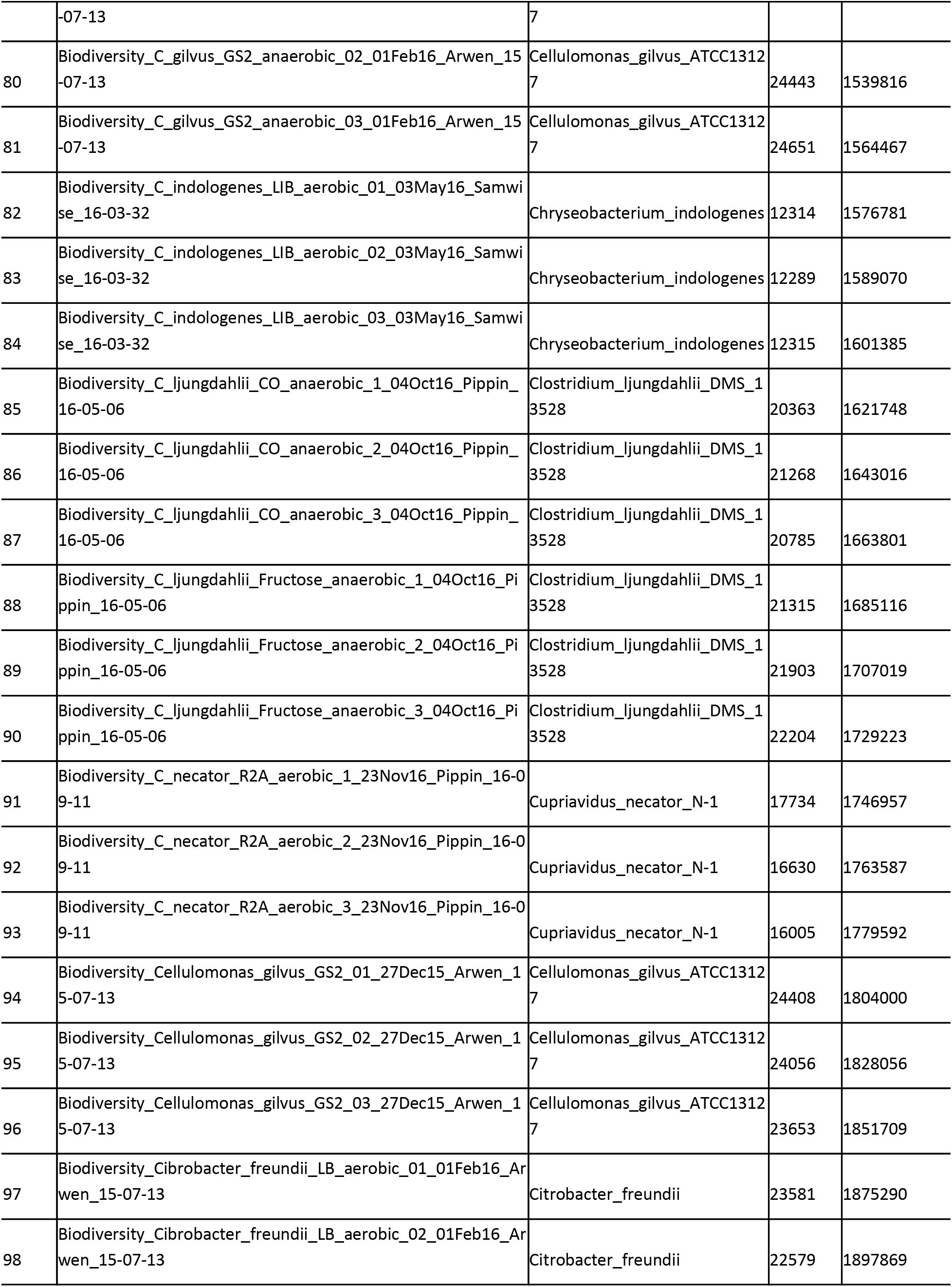
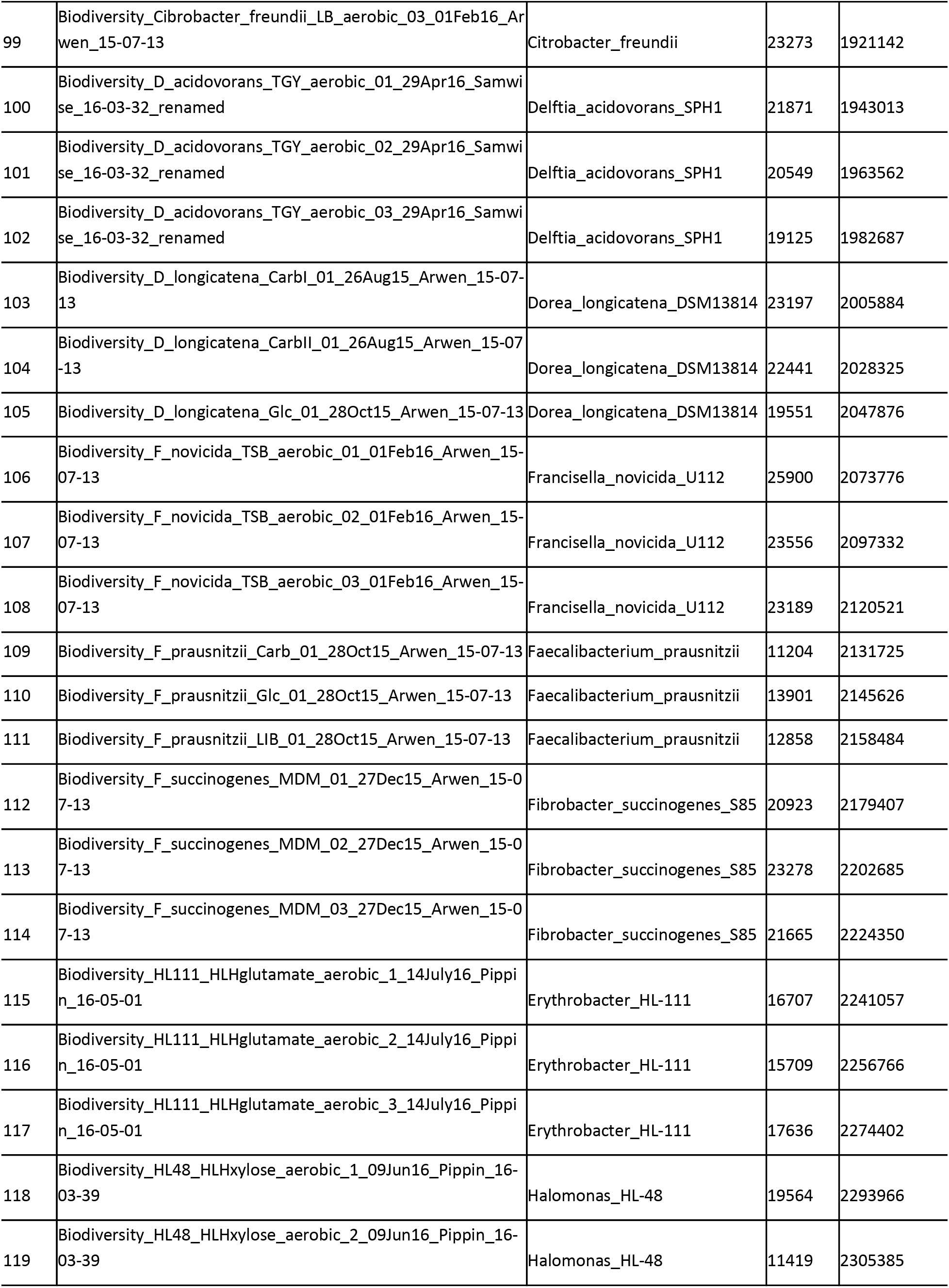
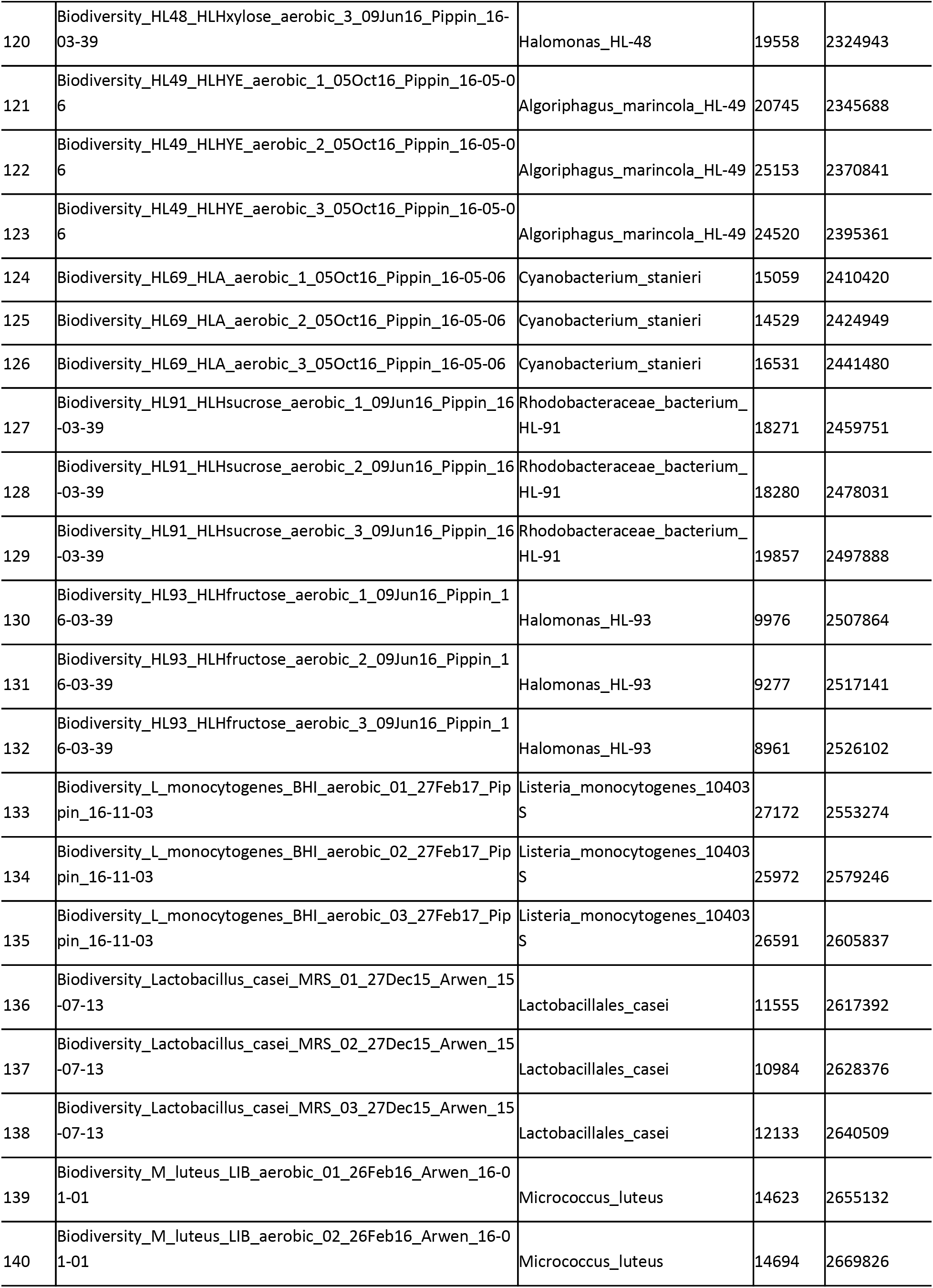
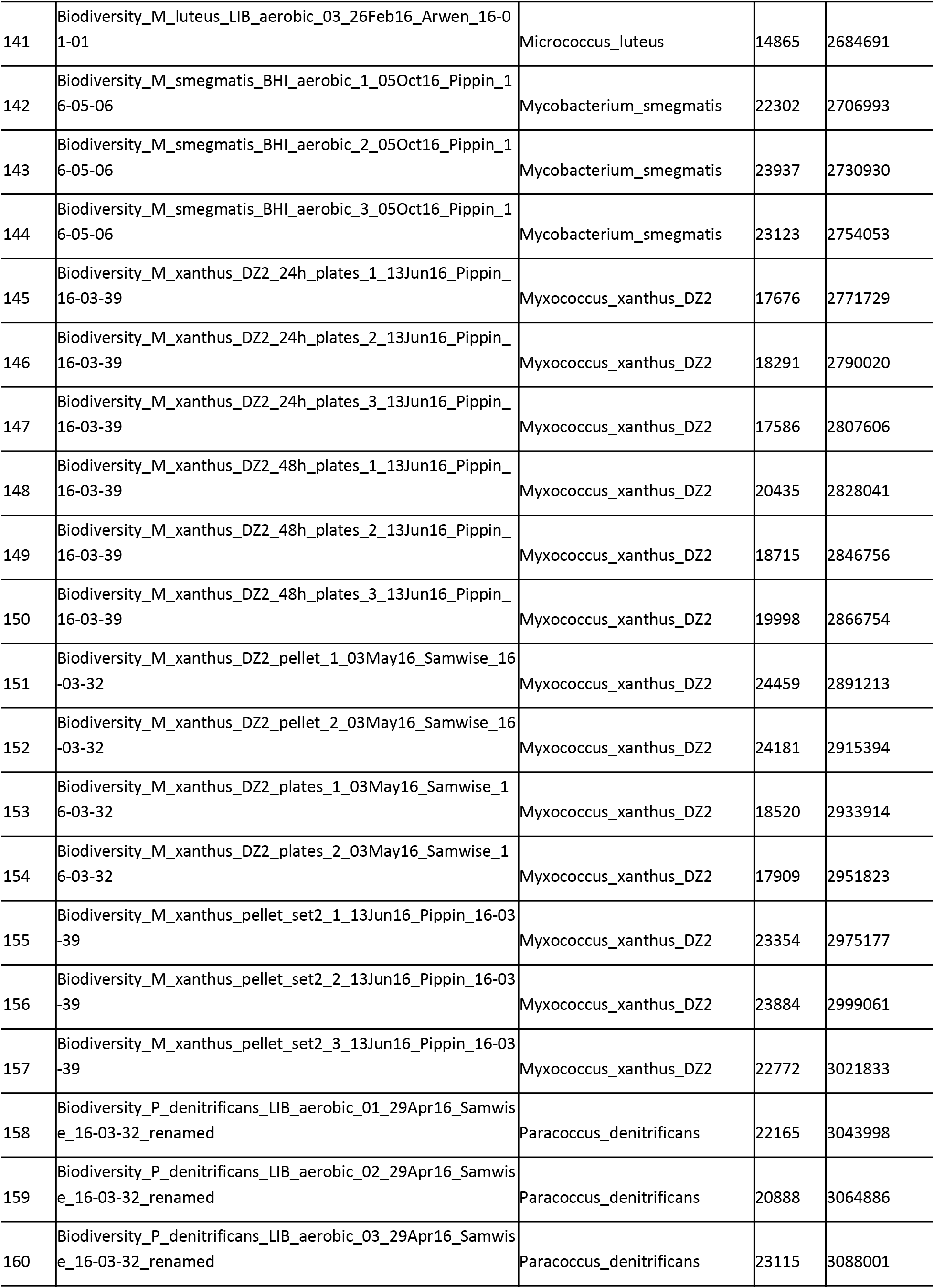
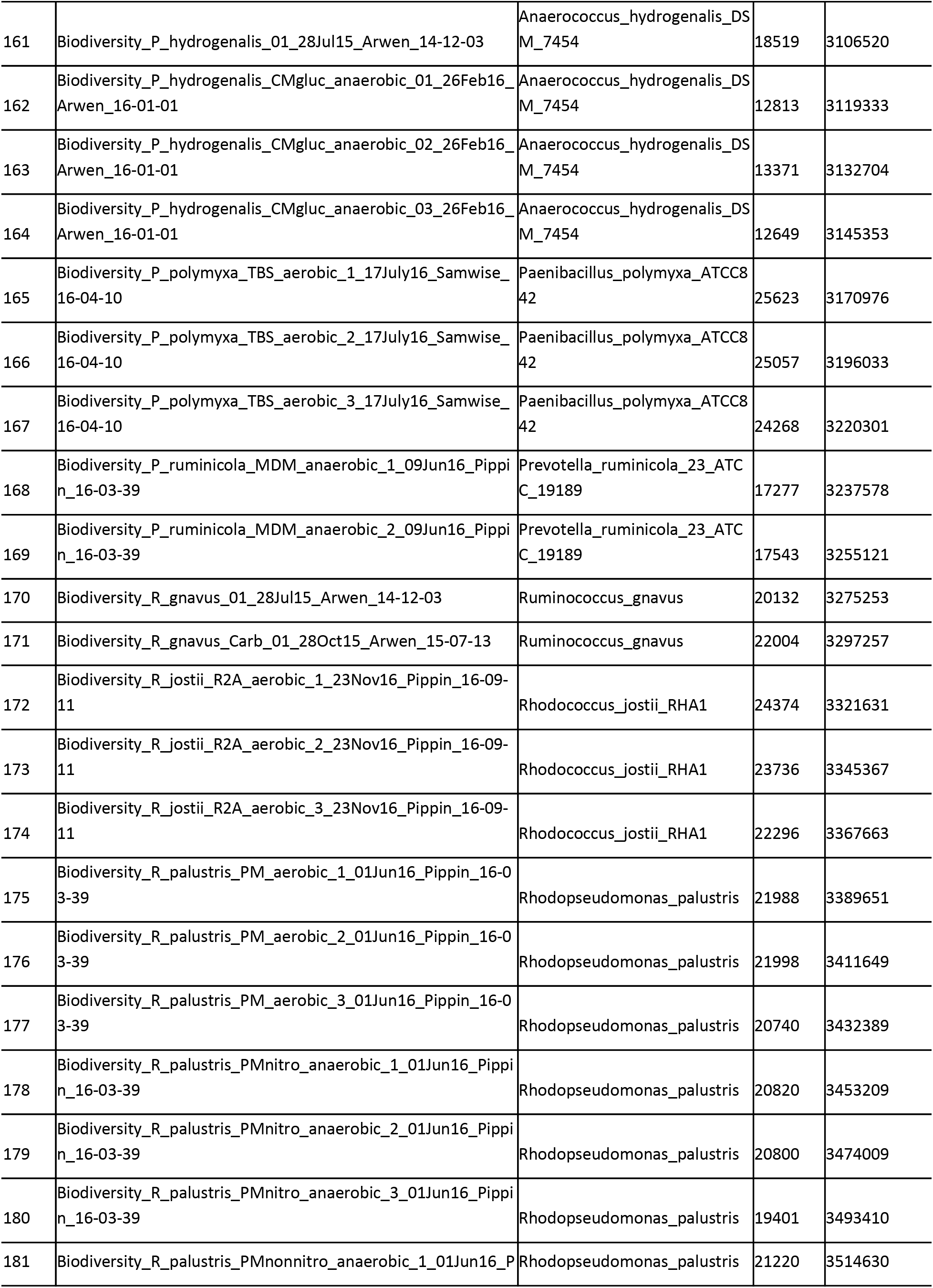
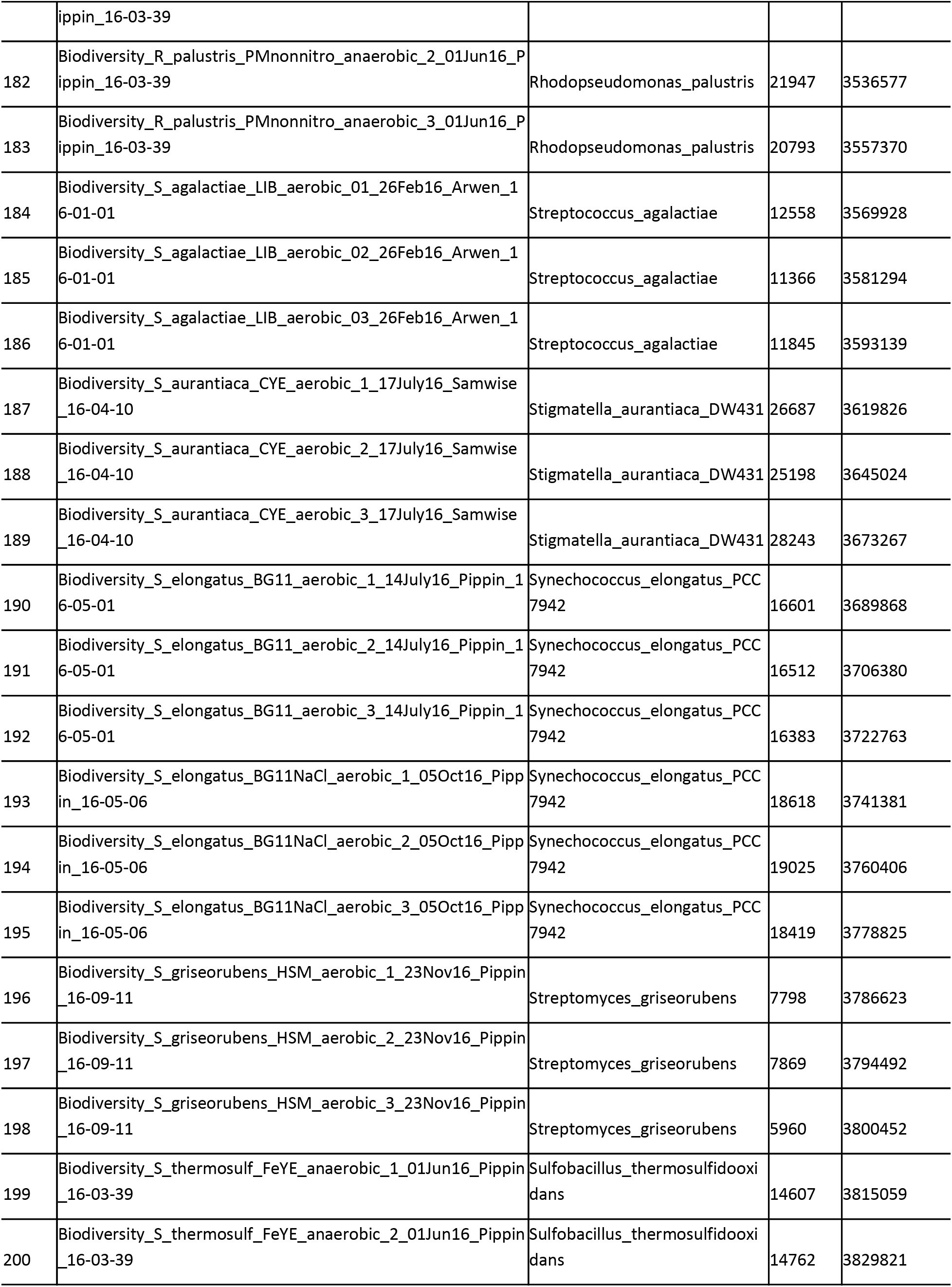
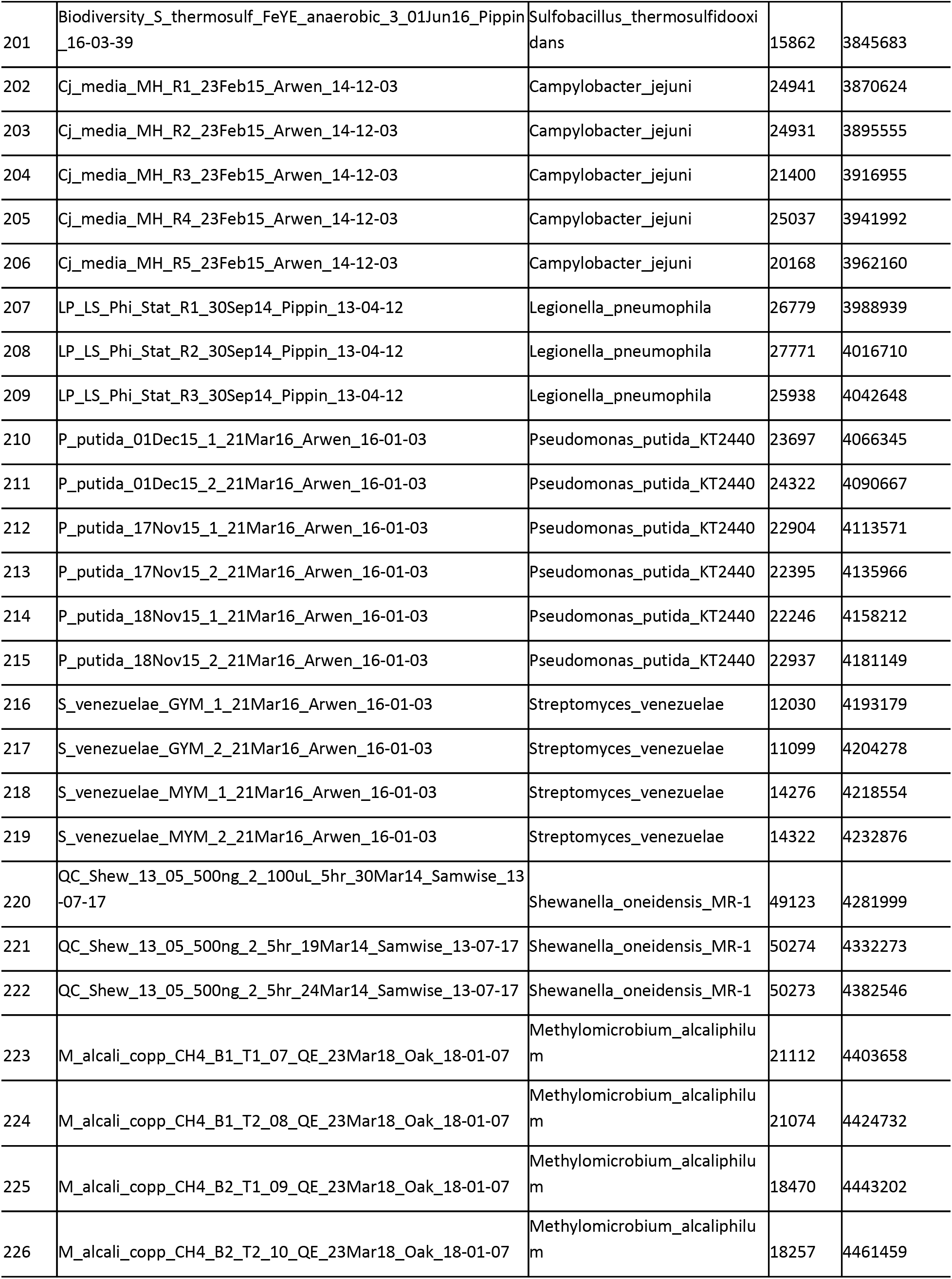
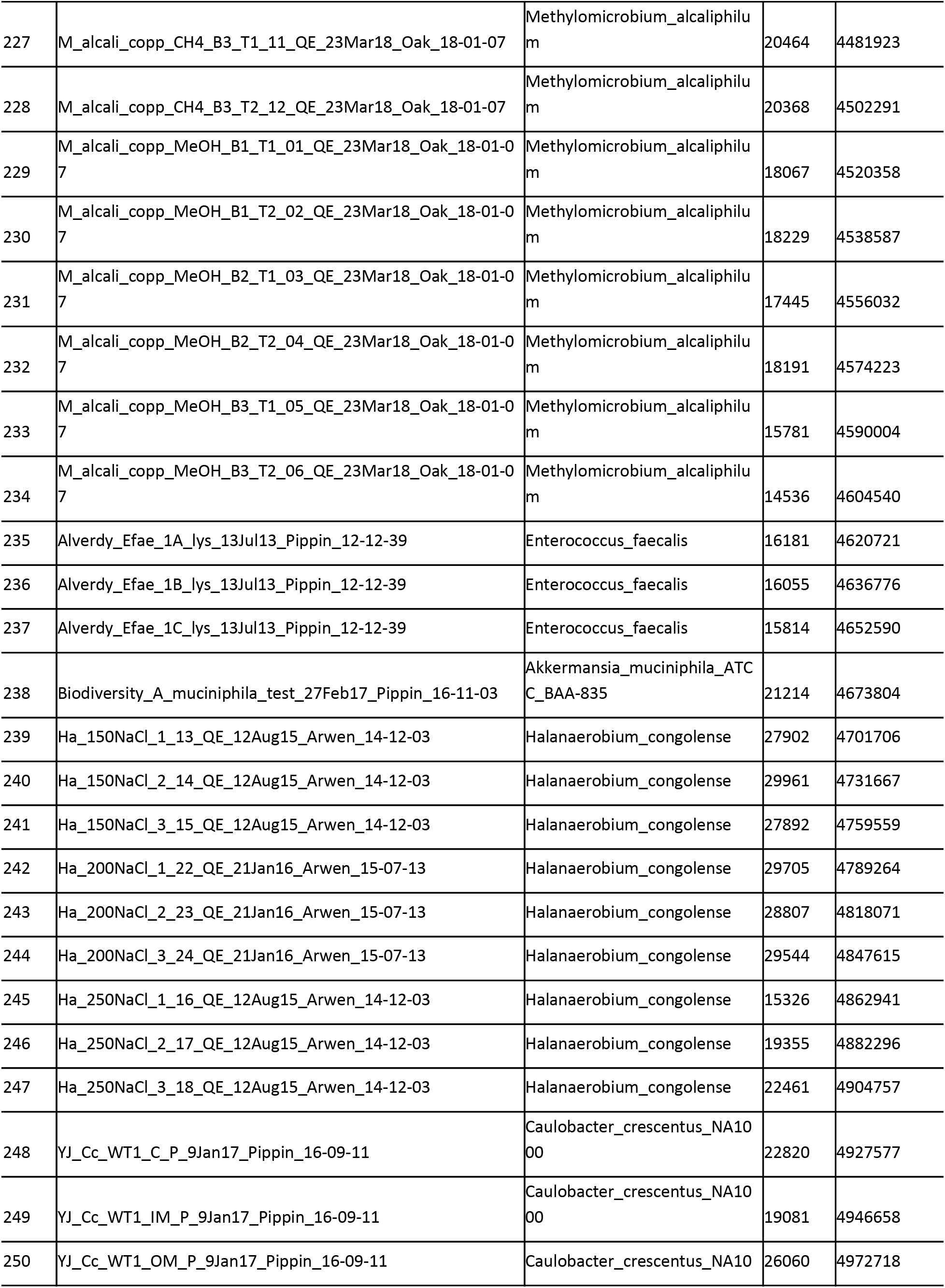
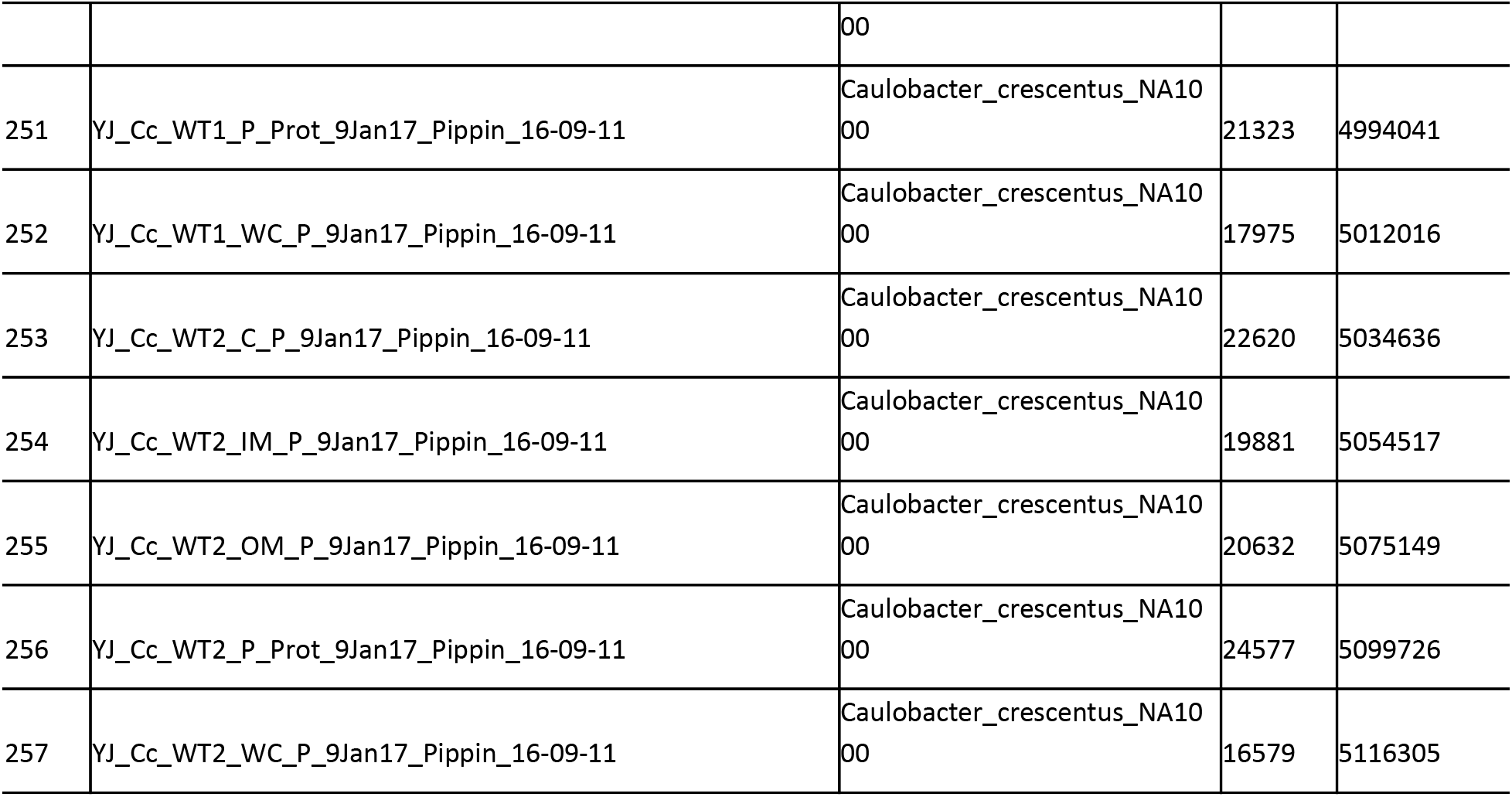
LC/MS data files for training and testing the Kaiko model.

**Supplementary Table 2.**
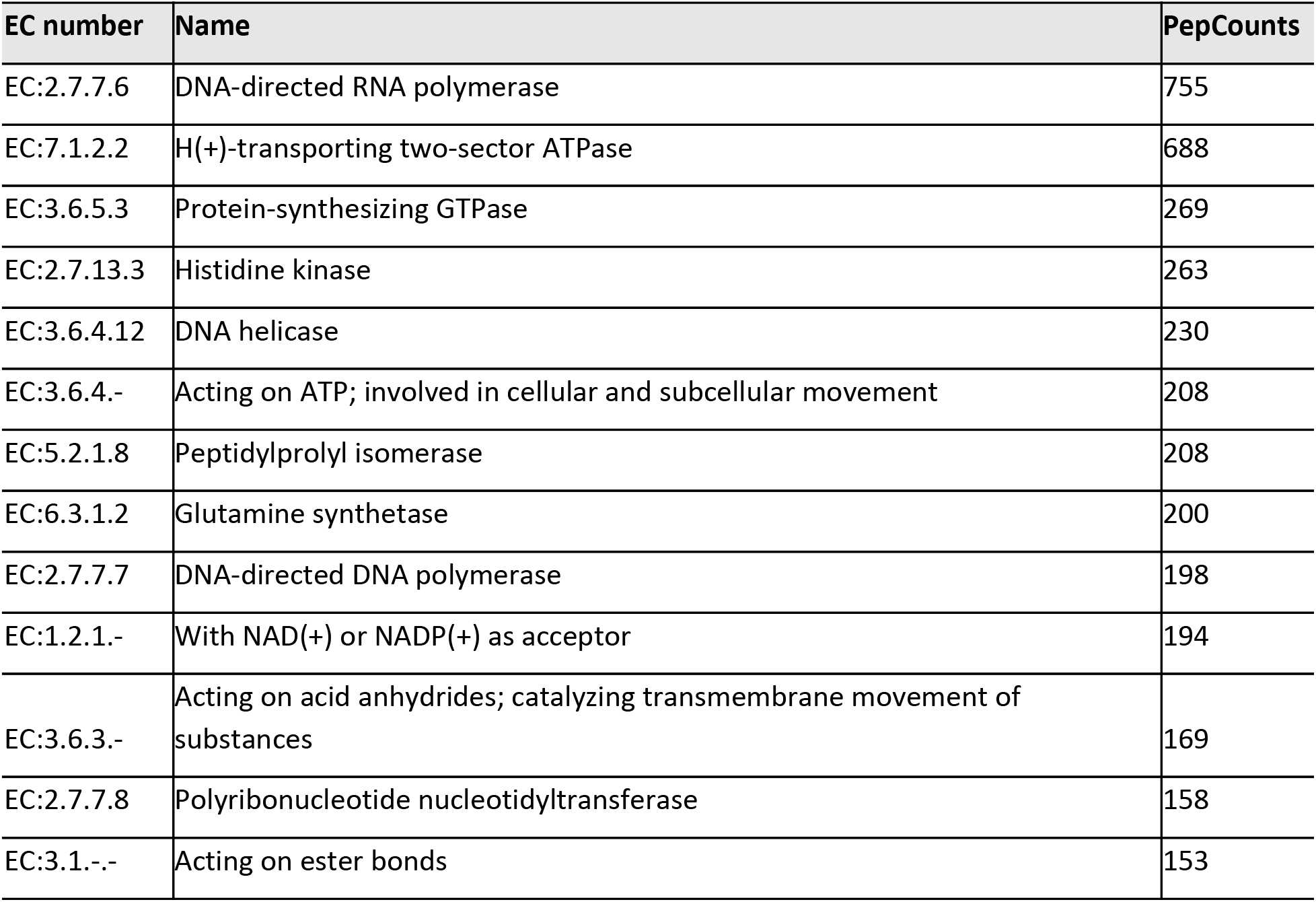
Top 20 of EC numbers most frequently matched from the unique peptides using Unipept 4.3 with the identified peptide sequences.

**Supplementary Table 3.**
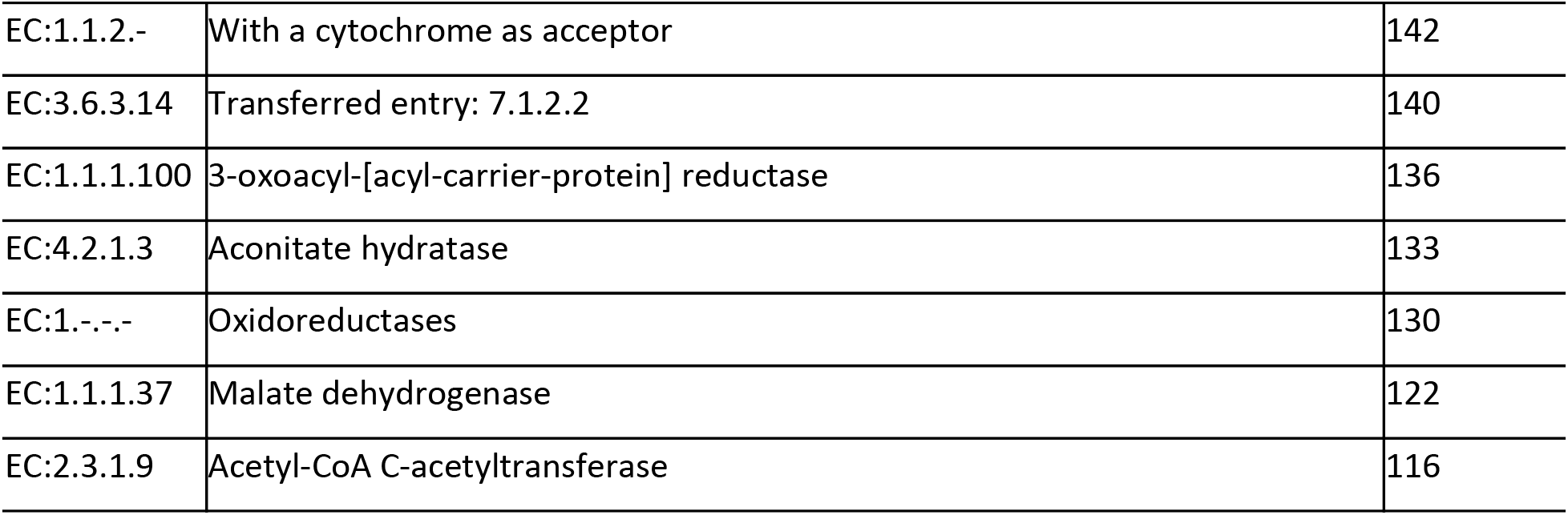
Functional distribution of the unique peptides across the phyla taxa. Each row and column represent the different EC numbers and phyla, respectively. The number in each cell indicates the number of unique peptides annotated by Unipept. This table was transformed from the original output file provided by Unipept. (It’s too big to add here so it will be attached as an Excel file)

